# Plant N-glycan breakdown by human gut *Bacteroides*

**DOI:** 10.1101/2022.04.07.487459

**Authors:** Lucy I. Crouch, Paulina A. Urbanowicz, Arnaud Baslé, Zhi-Peng Cai, Li Liu, Josef Voglmeir, Javier M. Melo Diaz, Samuel T. Benedict, Daniel I.R. Spencer, David N. Bolam

## Abstract

The major nutrients available to the human colonic microbiota are complex glycans derived from the diet. To degrade this highly variable mix of sugar structures, gut microbes have acquired a huge array of different carbohydrate-active enzymes (CAZymes), predominantly glycoside hydrolases, many of which have specificities that can be exploited for a range of different applications. Plant N-glycans are prevalent on proteins produced by plants and thus components of the diet, but the breakdown of these complex molecules by the gut microbiota has not been explored. Plant N-glycans are also well characterised allergens in pollen and some plant-based foods, and when plants are used in heterologous protein production for medical applications, the N-glycans present can pose a risk to therapeutic function and stability. Here we use a novel genome association approach for enzyme discovery to identify a breakdown pathway for plant complex N-glycans encoded by a gut *Bacteroides* species and biochemically characterise five CAZymes involved, including structures of the PNGase and GH92 α-mannosidase. These enzymes provide a toolbox for the modification of plant N-glycans for a range of potential applications. Furthermore, the keystone PNGase also has activity against insect-type N-glycans, which we discuss from the perspective of insects as a nutrient source.

## Introduction

Complex carbohydrates from a wide range of sources are the major nutrients available to the colonic microbiota. Degradation of these complex macromolecules by the microbiota is achieved through the massive expansion in genes encoding carbohydrate-active enzymes (CAZymes), with some species of gut microbe encoding >300 CAZyme genes from different families. *Bacteroides* species are particularly adept at glycan breakdown and typically organise the genes encoding the apparatus required for the breakdown of a particular glycan into discrete co-regulated loci (polysaccharide utilisation loci; PULs). A typical *Bacteroides* PUL comprises genes encoding CAZymes, the outer membrane glycan import system (SusC/D homologues), surface glycan binding proteins (SGBPs), and sensor-regulators (1). Discrete CAZyme gene clusters can also exist without the Sus or other PUL components, co-regulated with the core PUL apparatus located elsewhere on the genome. As a general rule, the more complex the substrate, the higher the number of CAZymes, CAZy families, and number of loci involved. For example, in *Bacteroides thetaiotaomicron*, the chemically simple substrate starch requires only one PUL, whereas breakdown of highly variable O-glycans induce the upregulation of 15 PULs(2).

N-glycans are common decorations of secreted proteins from almost all types of organisms and play important roles in protein stability and function(3). Although the core structures of plant and mammalian N-glycans are conserved, key differences exist in the types of sugar decorations and linkages. As a broad classification, mammalian complex N-glycans commonly have an α-1,6-fucose linked to the base GlcNAc whereas plants frequently have an α-1,3-fucose on this sugar and a β-1,2-xylose linked to the first mannose (Fig. 1). Insect N-glycans commonly have both α-1,3- and α-1,6-fucose linked to the core N-glycan. Plant complex N-glycans also differ from mammalian complex N-glycans in their antennae structures, which have β1,3-galactose and α1,4-fucose decorating the GlcNAcs (Fig. 1)(4), although there is significant variation in plant N-glycan structures depending on the species(5). Plant N-glycans are also of interest as they can be highly antigenic and induce allergic responses in mammals, causing both hayfever and food allergies(6-8). Furthermore when plants are used as hosts for heterologous protein production for medical applications and the N-glycans present on these therapeutic proteins can pose a risk to function and stability. Thus there is a need to be able to both characterise and modify plant N-glycans from different sources for a range of applications. Plant N-glycan specific CAZymes would be useful tools for this job, but currently there is a paucity of data describing enzymes that act specifically on plant N-glycan structures.

**Fig. 1.**
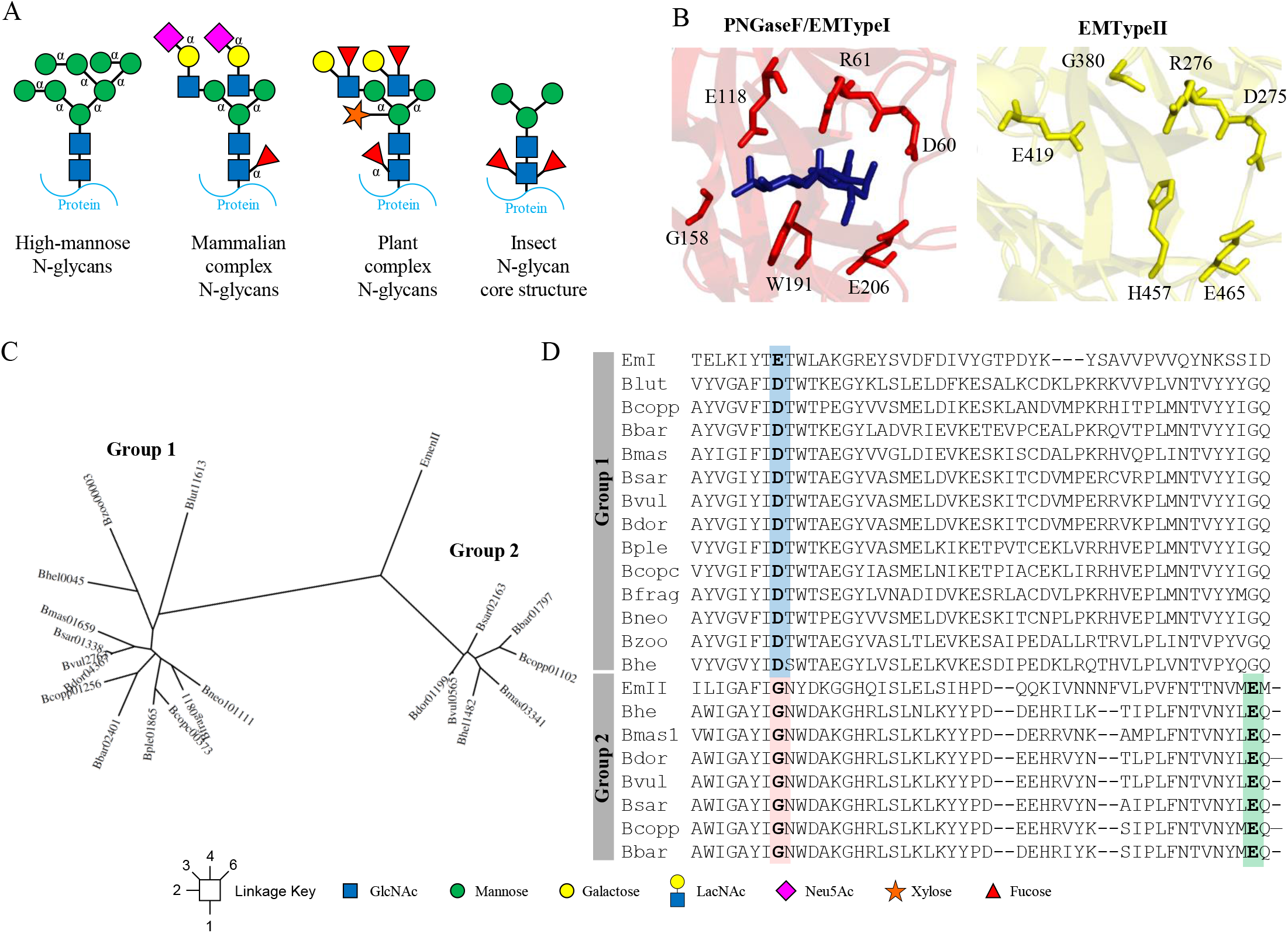
PNGase enzymes in species of Bacteroides. (A) Different types of N-glycans. High-mannose N-glycans (HMNGs) have mannose sugars decorating both arms usually to give a total of between 5 and 9 mannose sugars, dubbed Man5 and Man9, respectively, for example. HMNGs do not vary between different organisms, whereas complex N-glycans do have differences according to the source. In mammals, complex N-glycans have LacNAc disaccharides (Galβ1,4GlcNAc) attached to the mannose arms through a β1,2-linkage. The galactose sugars are typically decorated with sialic acids, but these can also decorate the antenna GlcNAc. Complex N-glycans can have addition antenna through a β1,4-linkage on the α1,3-mannose arm and a β1,6-linkage on the α1,6-mannose arm, to produce tri- and tetra-antennary structures, respectively. α1,6-fucose is a common decoration on the first core GlcNAc in mammals, but α1,3/4-linked fucose is also found to decorate the antenna GlcNAc. In contrast, Plant N-glycans typically have Lewis A epitopes as their antenna, a core α1,3-fucose, and a bisecting β-1,2-xylose. Insect N-glycan structures typically have both α1,3- and α1,6-fucose decorating the core GlcNAc. (B) The active sites of the two PNGases from *Elizabethkingia meningoseptica*. The key active site residues are shown as sticks and chitobiose is present in the PNGaseF/EMTypeI structure. (C) Phylogenetic tree of the PNGase enzymes from *Bacteroides* species, which broadly split into two groups. The members of Group 1 have quite variable identity between them, as low as 52 % in one instance, but generally between 67-99 %. The members of Group 2 have 75-97 % identity between them. (D) A sequence alignment to show residues that are key to the specificity of accommodating the α1,3-fucose typical of plant N-glycans. The residue blocking the α1,3-fucose in the Group I PNGases is highlighted in blue (E118 in PNGaseF/EMTypeI), the glycine replacing this residue in the Group 2 PNGases is highlighted in pink (G380 in EMTypeII), and the glutamic acid replacing the function of E118 is highlighted in green(E419 in EMTypeII).

Previous studies investigating carbohydrate-degradation systems in gut bacteria have typically used transcriptomic during growth on a specific glycan to identify the PUL or PULs involved in its breakdown(9, 10). These experiments require relatively large amounts of substrate for the bacteria to grow on, but for some glycans it may not be possible to isolate enough material, which means that discovery of enzymes that act on these substrates is currently limited. Plant N-glycans are a good example as although these molecules are common components of plant material and therefore widely consumed in the human diet, they are not easily available in the amounts required for transcriptomic studies (9). Here we describe a genome mining approach - “PULomics” - to look for the enzyme apparatus in *Bacteroides* species that degrade complex plant N-glycans. We relied on information about activity and specificity for particular enzyme families, assessed them for activity, and also extended this analysis to neighbouring putative CAZyme genes. Here we describe the biochemical characterisation of five CAzymes and two crystal structures to provide a degradation pathway for plant complex N-glycans encoded by the human gut microbiome. The study provides a toolbox of activities that will allow modification of plant and insect N-glycan structures for biotechnological and medical applications.

## Results

### Bioinformatics analysis shows two types of PNGase are present in *Bacteroides* species

There are currently two main classes of enzyme that remove N-glycans from glycoproteins; glycoside hydrolases (from either GH18 or GH85 families) and Peptide-N4-(N-acetyl-β-glucosaminyl)asparagine amidases (PNGases). The GH18 and GH85 enzymes hydrolyse the β1,4 glycosidic bond between the two core GlcNAc sugars, whereas PNGase family members cleave the linkage between the first GlcNAc and Asn of the protein/peptide. PNGaseF (or EMTypeI) from *Elizabethkingia meningoseptica* is widely used to remove mammalian-type N-glycans, which commonly have an α1,6-fucose attached to the first core GlcNAc (known as ‘Type I’ activity)(11). However, this enzyme is not able to accommodate N-glycan structures with α1,3-fucose attached to the core GlcNAc that are common decorations in plant and insect N-glycans(11). More recently, a ‘Type II’ PNGase also from *E. meningoseptica* was characterised that displayed additional activity towards N-glycans with a core α1,3-fucose (EMTypeII)(11). Structures of both PNGase enzymes exist and allowed structural insight into how these different α-fucose decorations are accommodated or blocked(11). In PNGaseF/EMTypeI, the active site residue Glu118 blocks where an α1,3-fucose may have potentially been accommodated, whereas in the EMTypeII structure, the equivalent residue is a Gly350, thus creating a pocket for this sugar (Fig. 1*B*). Glu418 in EMTypeII likely carries out the equivalent coordinating role to Glu118 in EMTypeI(11).

Recent work in *Bacteroides thetaiotaomicron* has described the degradation of mammalian-type biantennary complex and high-mannose N-glycans(9, 12). Both of these systems use a GH18 family member to remove the N-glycan from glycoproteins. *B. fragilis* has also been shown to degrade mammalian complex N-glycan structures(13) and a number of other *Bacteroides* species have the capacity to grow on glycoproteins with complex N-glycans(9). We wanted to further investigate the N-glycan degradation capacity of prominent gut *Bacteroides* species by exploring the prevalence and function of putative PNGases encoded by these prominent symbionts.

Analysis of The Integrated Microbial Genomes and Microbiomes system (IMG)(14) revealed the presence of putative PNGases in 13 species of *Bacteroides*, with 7 of these species having two genes each. These PNGase sequences clustered broadly into two groups according to sequence identity (Fig. 1*C*; Table 1). Sequence alignment included comparison to those from *E. meningoseptica* and revealed an interesting trend in terms of the possible substrate preferences of the two groups (Fig. 1*D*). Group 1 has aspartate residues in the equivalent position to Glu118 in EMTypeI, whereas Group 2 all had glycines at this position (Fig. 1*D*). This indicated that Group 1 and Group 2 *Bacteroides* PNGases may have similar substrate preferences to PNGaseF/EMTypeI and EMTypeII, respectively, in terms of their ability to accommodate a core α1,3-fucose. Structures for the Group 1 PNGases from *B. fragilis* and *B. vulgatus* are also available and confirm a similar positioning of the key catalytic residues relative to PNGaseF/EMTypeI (Fig. S1).

Another striking difference between the two groups is the presence of an N-terminal domain in most of the Group 2 protein sequences (all except *B. coprophilus*) and not in the group 1 sequences (Fig. S2). This N-terminal domain is present in EMTypeII, but not PNGaseF/EMTypeI, and has previously been dubbed the N-terminal bowl-like domain (NBL)(11). It has a unique structure and unknown function, but the expression of the catalytic domain alone without the NBL did not affect activity or specificity of EMTypeII (11).

A final noticeable difference between the two groups is a small insert between the two eight-stranded antiparallel β sheets in the sequences in Group 1 (Fig. S2). In the two available crystal structures of Group 1 PNGases from *B fragilis* and *B. vulgatus*, this translates into a small β-sheet twist adjacent to the active site (Fig. S1). Notably, PNGaseF/EMTypeI does not have this insert. Further analysis of the crystal structures of the PNGases from *B. fragilis* and *B. vulgatus* reveals that in both cases these enzymes crystallised in as dimers in the asymmetric unit with the twist being the major interaction and the active site was not blocked by this interaction (Fig. S1). This could be an indication of dimerization *in vivo* or an artefact of crystallography.

### The activity of Group 1 and 2 PNGases from *Bacteroides spp*

To explore the activity of the PNGases from *Bacteroides* species, a PNGase from each group (BF0811^PNGase^, a Group 1 enzyme from *B. fragilis* and B035DRAFT_03341^PNGase^, a Group 2 enzyme from *B. massiliensis*) was tested for activity against glycoprotein substrates displaying a range of different N-glycan types. These included fetuin, α1acid glycoprotein, RNaseB, and horseradish peroxidase (HRP), which have predominantly triantennary complex, biantennary complex, high mannose, and plant-type N-glycan decorations, respectively (Fig. 2). Commercially available PNGaseF/EMTypeI was also assayed for comparison. Released N-glycans were subsequently labelled with procainamide and analysed by liquid chromatography-fluorescence detection-electrospray-mass spectrometry (LC-FLD-ESI-MS). The *B. fragilis* PNGase from Group 1 (BF0811^PNGase^) displayed very similar activity to PNGaseF/EMTypeI with the removal of complex and high-mannose N-glycans, but not plant N-glycans (Fig. 2). In contrast, the *B. massiliensis* PNGase from Group 2 (B035DRAFT_03341^PNGase^) showed good activity towards plant-type N-glycans and limited activity towards the other substrates.

**Fig. 2.**
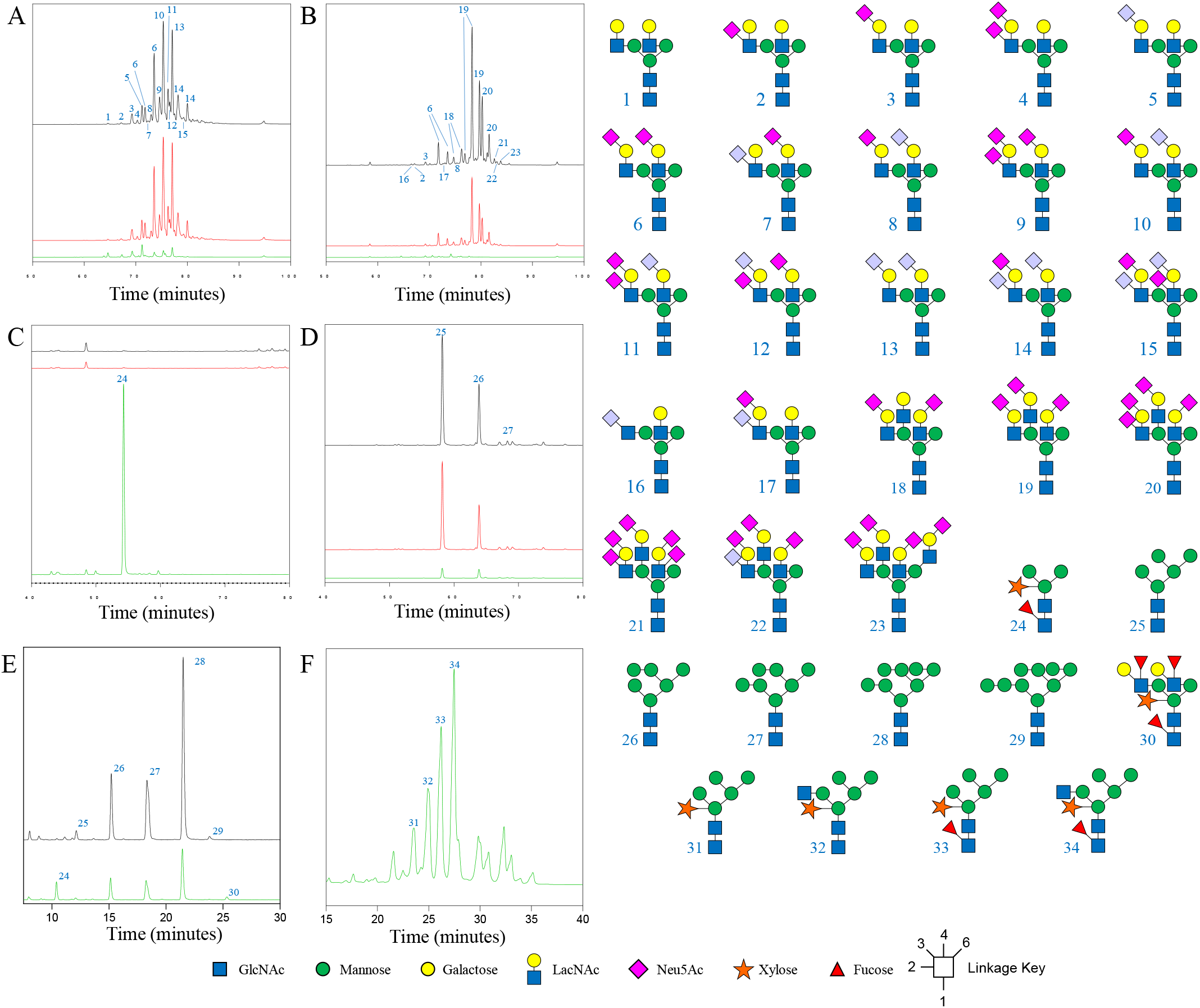
Activity of PNGase B035DRAFT_03341 from *Bacteroides massiliensis* against different substrates. (A) α_1_acid glycoprotein (B) Fetuin (C) Horseradish peroxidase (D) RNaseB (E) Soya protein (F) Papaya protein. B035DRAFT_03341 (green), PNGaseF (black), and BF0811 (red). The time window shown for the different chromatograms varies between the panels to provide clarity of the main peaks. The glycan products for A-E were labelled with procainamide and analysed by LC-FLD-ESI-MS. The glycan products for F were labelled with 2-AB and analysed by UPLC.

To further explore the substrate preference of B035DRAFT_03341^PNGase^, we used soya and papaya protein extracts as substrates (Fig. 2 and Figs. S3 & S4). The results demonstrated the ability of B035DRAFT_03341^PNGase^ to remove more decorated plant N-glycans, including those with high-mannose, hybrid, and complex antennary structures. For the soya protein, PNGaseF/EMTypeI was unable to remove structures with a core α1,3-fucose.

We also tested the activity of B035DRAFT_03341^PNGase^ against bee venom glycoprotein phospholipase A_2_, as this type of activity has previously been observed for EMTypeII(11). B035DRAFT_03341^PNGase^ was able to remove N-glycans from this insect glycoprotein. A decrease in molecular weight of phospholipase A_2_ was observed with the addition of B035DRAFT_03341^PNGase^ and there was also a decrease in intensity when staining for glycoproteins. This indicates the removal of N-glycan from this substrate (Fig. S5).

Notably, the *Bacteroides* PNGases are predicted to have a type I signal sequences, indicative of localisation to the periplasm (Table 2), which would suggest that deglycosylation occurs after the substrate has been imported across the outer membrane. This would be in contrast to GH18 directed cleavage of N-glycans which occurs at the cell surface in *B. thetaiotaomicron*(9, 12). One potential reason for this may be that the preferred PNGase substrates are glyco-peptides that are products of proteolytic digestion of glycoproteins, whereas GH18 enzymes deal with native glycoprotein. However, it is also possible that the signal sequences prediction is incorrect and the PNGases are localised to the outside, as has been seen previously(9).

### The structure of the *B. massiliensis* PNGase

To investigate the structural basis for specificity in the *Bacteroides* PNGase enzymes from Group 2, we solved the structure for B035DRAFT_03341^PNGase^ to 1.95 Å (Table S3). The structure consists of two domains: the catalytic domain and an NBL domain, which are linked through a flexible α-helical linker (Fig. 3A). The catalytic domain consists of two eight-stranded anti-parallel β-sheets, which is a consistent structural feature of PNGase enzymes described so far. The active site residues that are key for activity in EMTypeII are conserved in B035DRAFT_03341^PNGase^, with Gly388 occupying a critical position to allow the accommodation of α1,3-fucose (Fig. 3B). The available space for this fucose is particularly apparent in comparison to PNGaseF/EMTypeI (Fig. 3C). The α1,6-fucose points away from the active site so is not an issue in terms of blocking activity. The NBL domain has a very similar structure to the one in EMTypeII and is unique to these two proteins. They consist of 11 β-sheets connected by short α-helical regions and disordered loops (Fig. 3D).

**Fig. 3.**
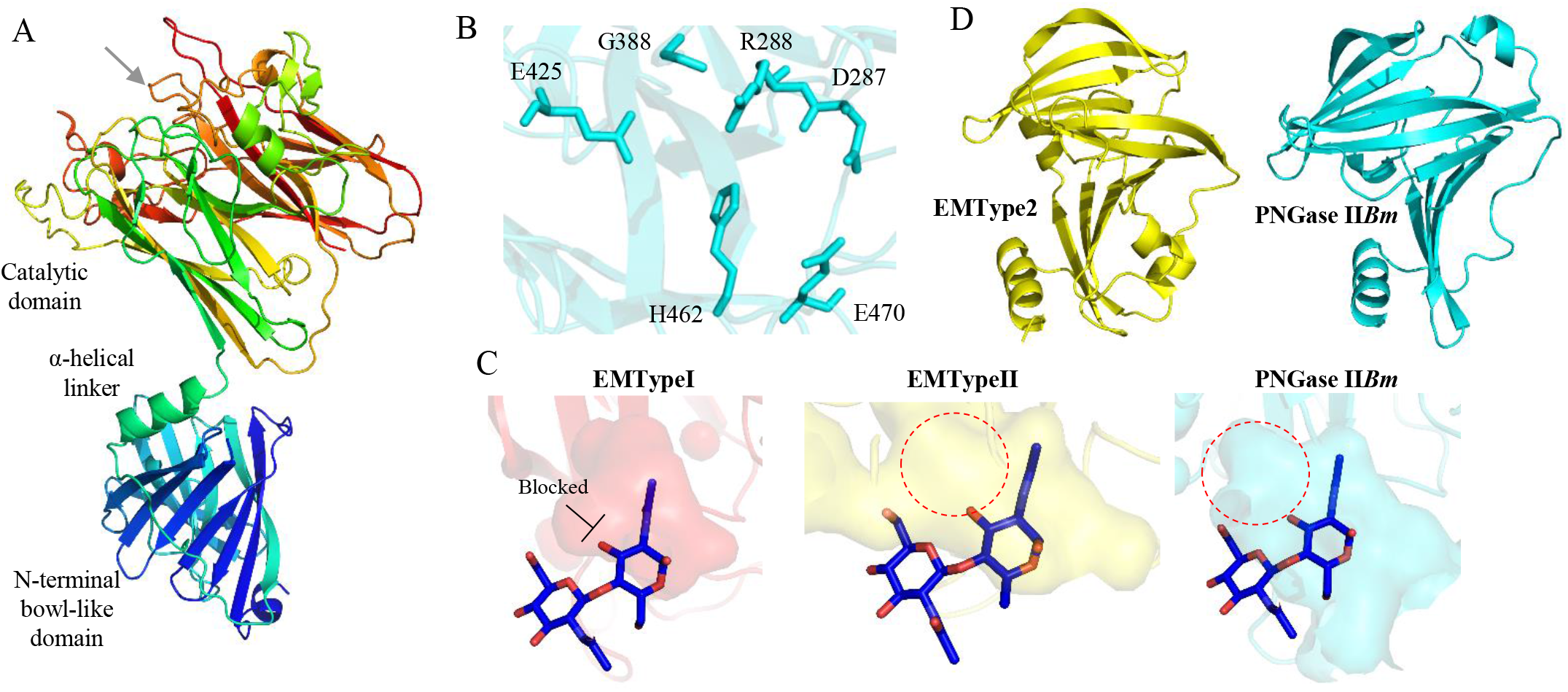
Structure of the Group 2 PNGase from *Bacteroides massiliensis*. (A) Structure of the Group 2 PNGase from *B. massiliensis* with two different domains. The protein is shown in a rainbow gradient with the N-and C-termini going from blue to red, respectively. The position of the active site is indicated (grey arrow). (B) The key catalytic residues of the active site. (C) The cavities and pockets are visualised for the active sites of three enzymes to show the space to accommodate α-1,3-fucose in EMTypeII and PNGase II*Bm* (red dotted circle) and also how it is not possible to accommodate this sugar in EMTypeI. (D) The two NBL domains from EMTypeII and PNGase II*Bm* to show their structural similarity.

### Plant N-glycan specific α-1,3-mannosidase

The presence of type II PNGase enzymes in prominent members of the human gut microbiota suggests that these microbes use plant N-glycans as a nutrient source. To identify the other enzymes required to fully degrade these glycans, we examined the putative CAZyme genes adjacent to the PNGase genes in ta number of different species of gut derived *Bacteroides*. The B035DRAFT_03341^PNGase^ gene is next to a putative GH92 (B035DRAFT_03340^GH92^) in *B. massiliensis*, the characterised members of which are all α-mannosidases (Fig. 4*A*). Notably, B035DRAFT_03340^GH92^ has a type II signal sequence suggesting it is membrane-associated, although whether the enzyme is localised to the cell surface or faces the periplasm in not known.

**Fig. 4.**
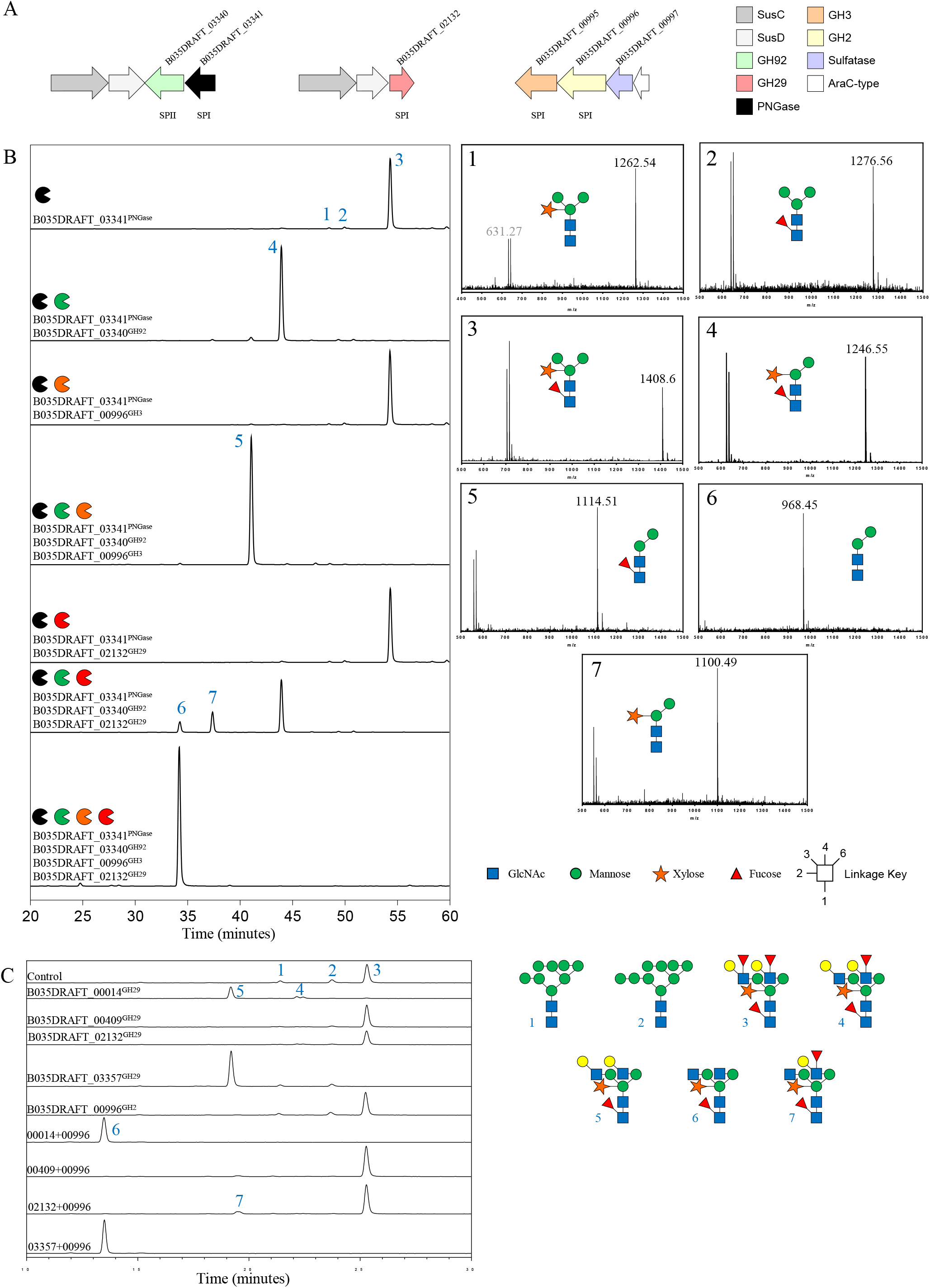
Enzymes highlighted from the functional association analysis. (A) Genes identified encoding the putative CAZymes highlighted by the functional association analysis carried out in search of enzymes involved in the degradation of plant N-glycans. (B) Horseradish peroxidase was incubated with different combinations of enzymes and the products were labelled with procainamide at the reducing end and analysed by LC-FLD-ESI-MS (left). The mass spectra of the different peaks are provided (right). (C) Soya bean N-glycans were incubated with GH29 fucosidases and B035DRAFT_00996^GH2^ galactosidase.

Incubation of this recombinant enzyme with α-mannobiose of varying linkages showed activity against the α-1,3 substrate only (Fig. S6). B035DRAFT_03340^GH92^ was then assayed against HRP, which has simple plant-type N-glycan decorations (no antenna decorations; Fig. S6). B035DRAFT_03340^GH92^ was able to remove the α-1,3-mannose from plant N-glycan heptasaccharide once it was removed from the protein and also whilst the glycan was still attached to the protein (Fig. 4*B &* Fig S6).

To further study the specificity of B035DRAFT_03340^GH92^ it was compared to a GH92 that has specificity towards the α-1,3-mannose linkages in high mannose N-glycans (BT3991^GH92^ from *B. thetaiotaomicron*)(12). BT3991^GH92^ was unable to remove mannose from HRP either with or without the removal of the N-glycan from the protein, whereas B035DRAFT_03340^GH92^ could cleave the α-1,3-mannose from both free HRP glycan or while the N-glycan was attached to protein (Fig. S4). The lack of activity seen for BT3991^GH92^ is likely due to the plant-specific β-1,2-xylose causing steric hindrance within the active site, whereas B035DRAFT_03340^GH92^ is able to accommodate this decoration.

Close homologues of B035DRAFT_03340^GH92^ are predicted to be in all the *Bacteroides* species that have a Group 2 PNGase encoded in the genome. These homologues had identity between 77 and 96 % (Table S4) and are present adjacent to the gene for the Group 2 PNGase for *B. massiliensis, B. vulgatus*, and *B. helcogenes*, but not in the case of *B. dorei, B. sartorii, B. coprophilus*, or *B barnesiae*. An exception is *B. neonati* that also has a homologue of B035DRAFT_03340^GH92^ with 73 % identity, but no putative Group 2 PNGase gene and, unexpectedly, this GH92 gene is adjacent to the putative Group 1 PNGase in *B. neonati* (Fig. S7).

### The structure of the GH92 α-1,3-mannosidase able to target plant N-glycans

To investigate the structural basis for the unusual specificity displayed by B035DRAFT_03340^GH92^ we determined the crystal structure of the enzyme to 1.43 Å (Table S3). The enzyme consists of two domains: an N-terminal β-sandwich domain composed of 16 antiparallel β-strands domain and an (α/α)_6_-barrel catalytic domain. These two domains are pinned together by two α-helices, previously dubbed Helix 1 and 2 (Fig 5*A*)(15). The secondary structures of the seven GH92 enzymes with known structures also have these three features: the N-terminal domain, the catalytic domain, and the α-helical linker. The density for several metal ions was observed bound to the protein surface, which were modelled as Na from the crystallisation conditions, except for a Ca near the active site, as this metal has been shown to be key for GH92 activity.

**Fig. 5.**
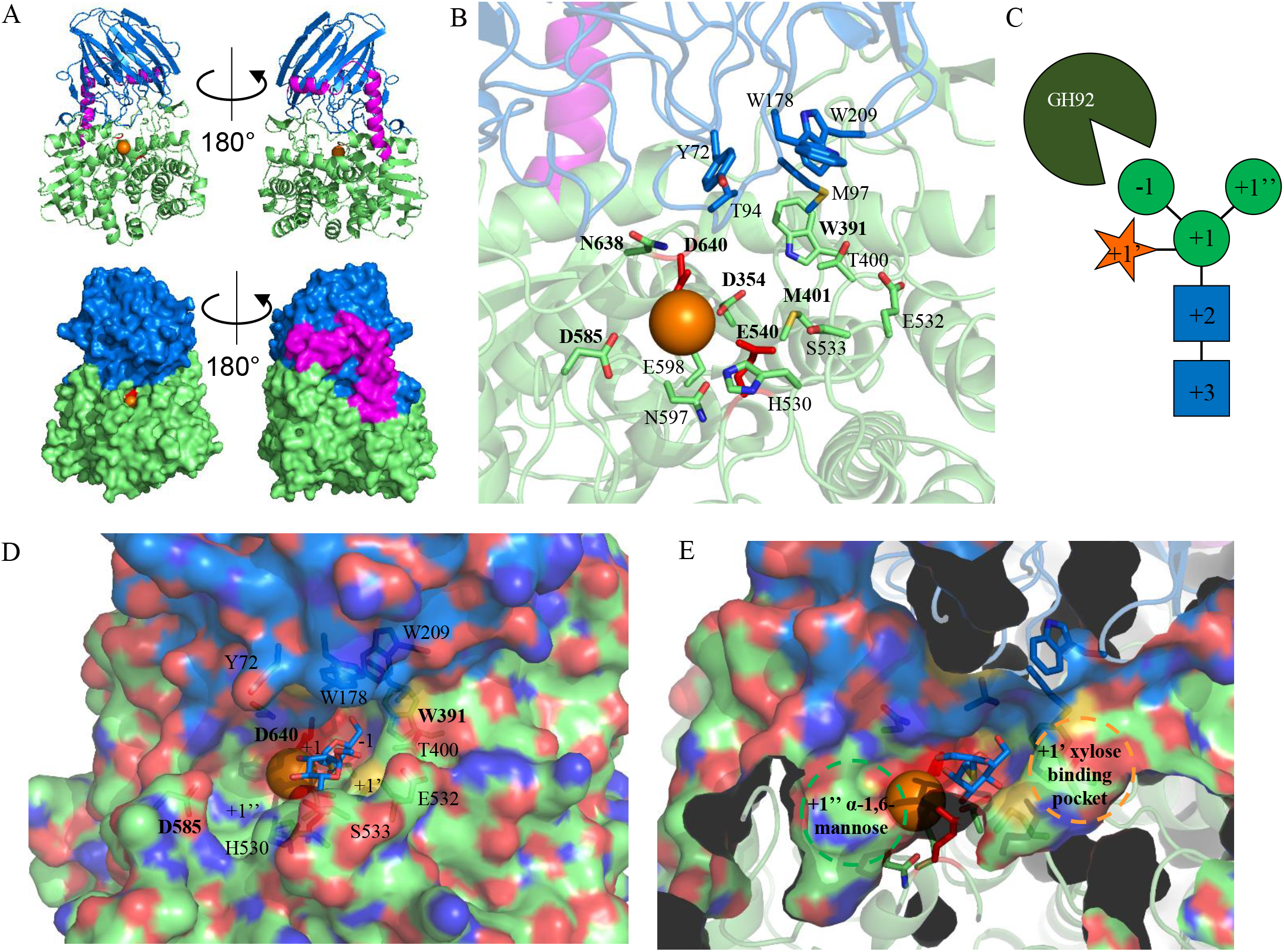
Crystal structures of the *α*-1, 3-mannosidase B035DRAFT_03340^GH92^. (A) Structures of B035DRAFT_03340^GH92^ shown in cartoon and surface, top and bottom, respectively. The N-terminal β-sandwich domain, the two connecting helices, and the C-terminal (α/α)_6_-barrel are shown in marine, magenta, and lime, respectively. The metal ion is shown in orange and catalytic residues are shown in red. (B) Details of the active site with likely important residues shown as sticks. Those labelled in bold are conserved throughout the GH92 family and those not in bold are unique to this enzyme. (C) Diagram showing the sugar subsites for this enzyme. (D) Surface representation of the active site with partial transparency so the residues in sticks can also be viewed. Thiomannobiosidefrom BT3990 (PDB2WW1) has been overlaid to show the approximate -1 and +1 subsites. The likely +1’ and +1’’ are also labelled. (E) This is the same as panel D, but showing a cross section through the front of the enzyme to show the extent of the binding pockets for the +1’ and +1’’ subsites.

The active site of B035DRAFT_03340^GH92^ is comprised of residues originating from both the N- and C-terminal domains, which is a common feature of the other GH92 structures (Fig 5*B*). The residues that are conserved within the catalytic site throughout all GH92 enzymes with structures derive from the C-terminal domain. Those residues that vary come from both domains and are the drivers of specificity by interacting with the +1 subsite sugar and beyond. To explore the active site of B035DRAFT_03340^GH92^, we overlaid the nonhydrolysable substrate mimic thiomannobioside from the structure of BT3990 (PDB 2WW1) (Fig 5*D*). From this we could speculate about four subsites, including the -1 α-1,3-mannose, +1 core mannose, +1’ xylose and the +1’’ α-1,6-mannose. The environment of the -1 mannose is identical to that described for other GH92 enzymes structures(15-17). The +1 mannose is likely coordinated by at least three residues: H530, S533, and E532 which come up from underneath the mannose. Overhanging this subsite (and possibly the +2 GlcNAc position also) are three hydrophobic residues Y72, W178, and W209, which is suggestive of π-stacking of the sugars. However, these aromatics are more distant than equivalent residues seen in other GH92 structures(15). The +1 subsite interactions allows space either side of the +1 mannose for the +1’’ mannose and +1’ xylose. There are clear pockets for these sugars in the B035DRAFT_03340^GH92^ structure (Fig 5*E*).

The eight GH92 structures already available include five α-1,2-mannosidases, an α-1,3-mannosidase, an α-1,4-mannosidase, and a mannose-α-1,4-PO_4_-mannose mannosidase(15-19). Previous comparison of the α-1,2-mannosidase structures revealed three residues coordinating the mannose at the +1 subsite that drive specificity for α-1,2-linkages. These are a Trp from the N-terminal domain and a Glu and His from the C-terminal domain and these are also predicted through sequence alignments to be present in other GH92 α-1,2-mannosidases. SP2145 from *Streptococcus pneumoniae* PDB 5SW1 was crystallised with a mannose in the +1 subsite and demonstrates these interactions (Fig S8). In an attempt to highlight if there were any similar conserved motifs present for GH92 α-1,3-mannosidases, we compared the structures of B035DRAFT_03340^GH92^ with BT3130 (PDB 6F8Z; Fig S8). This comparison saw no conservation in the active site residues associates with the +1 subsite. However, the residues contributed from the N-terminal domain were tryptophans, like those seen in the α-1,2-mannosidases, but the location and orientation differed (Fig. S8). Notably, in B035DRAFT_03340^GH92^ this tryptophan +1 subsite “lid” is much further away from where the glycan would sit than in other GH92 structures. This lid would possibly reach down further if substrate was present.

We carried out phylogenetic analysis of the GH92 enzymes that had been characterised to see if they would cluster according to their activities (Fig S9). This was successful in that α-1,2-mannosidases and α-1,3-mannosidases clustered together. The sequences were predominantly derived from *B. thetaiotaomicron*, so this may not be a completely reliable method of predicting specificities.

### Gene association analysis to identify additional plant N-glycan degrading enzymes

The characterisation of PNGase homologues from gut *Bacteroides* species revealed there are likely two different PNGase-like activities encoded by these microbes; Group 1 targeting mammalian N-glycans and Group 2 targeting plant N-glycan structures. We were also able to identify an α-1,3-mannosidase with specificity towards plant-type N-glycans by characterising the product of a GH92 gene associated with the Group 2 PNGase gene in *B. massiliensis*. Genes in the same locus likely have functional associations and this is common in carbohydrate degradative systems in Bacteroidetes. We therefore expanded this concept to identify other putative plant N-glycan targeting CAZymes from *Bacteroides* species.

The group 2 PNGases from *B. dorei, B. barnsiae* and *B. coprophilus* are all orphan genes (i.e. no obvious adjacent genes) and the PNGase from *B. sartorii* only neighbours a susC/D pair, however the Group 2 PNGase genes from *B. helcogenes* and *B. vulgatus* all look to be a part of more extensive loci (Fig. S7). The neighbouring ORFs included putative SusC/D pairs, GH29, GH3, GH130, and additional GH92 enzymes (Fig. S7). Using this initial survey, a network of possible functionally related ORFs was built for all the *Bacteroides* species with putative Group 2 PNGase enzymes (Fig. S7). Using this approach, we were able to highlight CAZymes with potential specificity towards plant N-glycans. For *B. massiliensis*, these CAZymes were located in two further putative loci (Fig. 4*A*). One locus consists of a susC/D pair and a putative GH29 and the second has a GH3, a GH2, a sulfatase, and an AraC-type regulator. The activities of these *B. massiliensis* CAZymes were then explored.

### B035DRAFT_00995^GH3^ is a β-xylosidase acting on plant N-glycans

B035DRAFT_00995^GH3^ was screened against a variety of pNP substrates and found to be active against pNP-β-xylose. Using this information, we then incubated this enzyme with the plant-type heptasaccharide released by B035DRAFT_03341^PNGase^ (Fig. 4*B*). This was unsuccessful at removing the bisecting β-1,2-xylose. However, when the reaction was carried out also in the presence of B035DRAFT_03340^GH92^ to remove the α-1,3-mannose, B035DRAFT_00995^GH3^ was able to remove this xylose (Fig. 4*B &* Fig. S10). B035DRAFT_00995^GH3^ has a Type I signal peptide, so is likely localised to the periplasm (Table 2).

B035DRAFT_00995^GH3^ activity was also assessed against β-1,4-xylobiose (Fig. S10). No activity was observed, which indicates specificity for this enzyme towards β1,2 linkages in plant N-glycans. For comparison, a previously characterised GH3 β-xylosidase from *B. ovatus* (BACOVA_03419) that is involved in the degradation of plant cell wall xylans, could hydrolyse β-1,4-xylobiose (Fig. S10).

Putative GH3 enzymes were identified for 5 out of the 7 *Bacteroides* species with Group 2 PNGase enzymes. Homologues of B035DRAFT_00995^GH3^ in *B. coprophilus, B. barnsiae, B. vulgatus*, and *B. helogenes* have 66, 67, 75 and 76 % identity, respectively (Table S5). No obvious equivalent B035DRAFT_00995^GH3^ homologues could be identified in *B. dorei* or *B. sartorii* using the functional association analysis.

### B035DRAFT_02132^GH29^ is an α-1,3-fucosidase specific to the core decorations of plant N-glycans

The fucose decorating the core GlcNAc of a plant N-glycan is through an α-1,3-linkage, in contrast to the α-1,6-linkage of mammalian-derived N-glycans. Another enzyme identified through the functional association analysis was B035DRAFT_02132^GH29^, which is predicted to be localised to the periplasm (Table S2). GH29 family members typically have exo α-1,3/4-fucosidase activities, so B035DRAFT_02132^GH29^ was screened against a variety of fucose-containing glycans (Fig. S11). B035DRAFT_02132^GH29^ was found to only hydrolyse the α-1,3-fucose from Lewis X trisaccharide to completion overnight, which is the glycan most similar to the core of a plant N-glycan out of the defined oligosaccharides that were tested. As a comparison it was only partially active against the α-1,3-fucose from 3-fucosyllactose, which confirms a specificity for GlcNAc over Glc in the +1 subsite. Furthermore, B035DRAFT_02132^GH29^ was not able to remove the α-1,4-fucose from Lewis A, which indicates that it does not target the antennary structures of plant N-glycans (Fig. S11).

When B035DRAFT_02132^GH29^ was tested against plant N-glycan heptasaccharide released by B035DRAFT_03341^PNGase^, no core fucose removal was observed (Fig. 4*B*). However, partial removal of the core α-1,3-fucose was seen once the α-1,3-mannose had been removed by B035DRAFT_03340^GH92^ and full removal of the α-1,3-fucose by the GH29 was made possible after removal of the β-1,2-xylose by B035DRAFT_00995^GH3^. These observations provide insights into the likely plant N-glycan degradation pathway in *B. massiliensis* (Fig. 6). Homologues of these enzymes were in five out of the seven *Bacteroides* species with TypeII PNGase enzymes (Table S6). There was no obvious homologue in *B. vulgatus* and *B. dorei* and there were also no obvious homologues in species without TypeII PNGases.

**Fig. 6.**
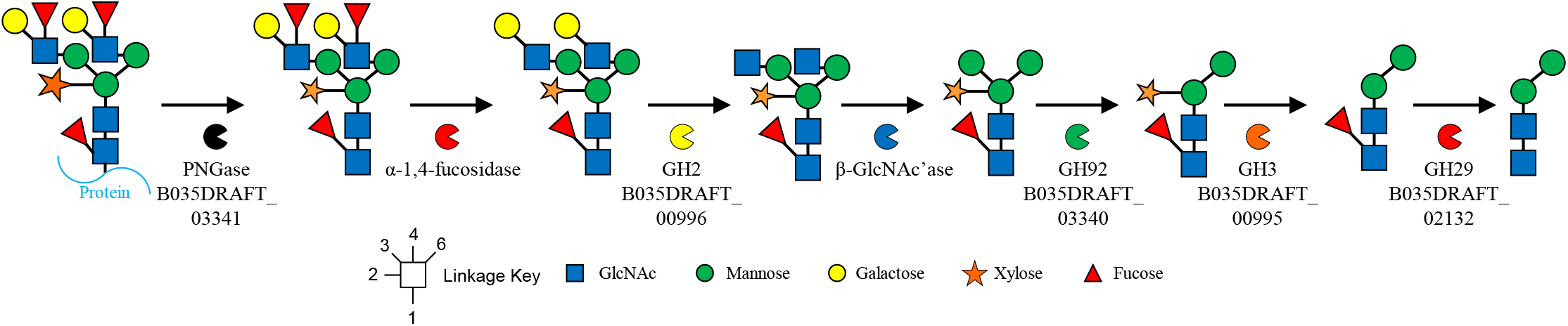
Model of the degradation of plant N-glycans by *Bacteroides massiliensis*. This summarises the order in which *B. massiliensis likely* degrades plant N-glycans.

Fucosidases from different sources have previously been shown to act on the α-1,3-linkage core linkage. Most notably, a GH29 from *E. meningoseptica*, cFase I, can act on the core α-1,3-fucose even when the plant N-glycan has antennary decoration(20). Another GH29 from *Arabidopsis thaliana*, AtFUC1, was also able to act on the α-1,3-linkage, but only when the glycan was reduced down to an α-1,3-fucose linked to chitobiose. A knockout of the AtFUC1 gene lead to an accumulation of this trisaccharide in the plant confirming the enzymes specificity towards the core fucose of plant N-glycans(21).

### Comparison between plant N-glycan degradation pathways in *B. massiliensis* and a bacterial phytopathogen

*Xanthomonas campestris* pv. *campestris* causes black rot disease in *Brassica* plant species and in a previous study a set of genes upregulated in the presence of GlcNAc was explored in terms of plant N-glycan degradation(22). These genes were predicted to be putative CAZymes from a range of families and their subsequent characterisation revealed some comparable observations to the work described here. Firstly, a GH92 (NixK) was able to remove the α-1,3-mannose from a plant N-glycan heptasacchairde, akin to what was observed here for B035DRAFT_03340^GH92^ (42 % identity between these two enzymes). Furthermore, without the removal of this mannose, the activity of other enzymes was blocked, as observed in the *B. massiliensis* system.

Plant N-glycan β-1,2-xylosidase activity was also observed with a GH3 family member (NixI) from *X. campestris*. NixI has a low identity to B035DRAFT_00995^GH3^ of 33 %, but the specificity of acting after the removal of the α-1,3-mannose is the same. In terms of core α-1,3-fucosidase activity, a GH29 (NixE) could remove this sugar, but only when all mannose sugars had been removed, which is not the case for B035DRAFT_02132^GH29^. It is worth noting that the substrate used in this study was a glycopeptide produced from trypsin degradation of avidine produced in corn and not a free N-glycan, which may influence the activities observed.

There was no endo-acting enzyme activity characterised for the *X. campestris* system, although a GH18 was present in the GlcNAc-activated locus. The GH18 is a likely candidate for removal of the N-glycan in *X. campestris*, unlike *B. massiliensis* which employs a PNGase.

### B035DRAFT_00996^GH2^ is a β-1,3-galactosidase specific to the antenna decorations of complex plant N-glycans

The final CAZyme identified in *B. massiliensis* using the functional association analysis was a GH2, B035DRAFT_00996^GH2^, which was screened against a variety of pNP substrates and found to be active against pNP-β-galactose. This enzyme was initially screened against defined oligosaccharides to determine its specificity (Fig. S12). B035DRAFT_00996^GH2^ only had activity towards β1,3-linked galactose when GlcNAc was in the +1 position (Lacto-N-biose). It could also act on Lacto-N-tetraose, which has the same linkage and +1 sugar. It was unable to hydrolyse LacNAc or Galβ1,3Glc, which demonstrates the specificity towards the β1,3-linkage and a requirement for the N-acetyl group of the +1 GlcNAc, respectively. Furthermore, partial activity was observed towards Galβ1,3GalNAcβ1,3Galβ1,4Glc, which emphasises the importance of the N-acetyl group in the +1 sugar with some influence also coming from the C4 hydroxyl of the +1 sugar either being axial or equatorial (Gal or Glc, respectively). These results show that B035DRAFT_00996^GH2^ has specificity towards the linkage and +1 GlcNAc sugar found on the antenna of complex plant N-glycans.

Activity was also tested against Lewis A trisaccharide, which is the epitope of the antenna structure present on plant N-glycans (Fig. 1*A &* Fig. S12). B035DRAFT_00996^GH2^ was unable to remove the galactose in this case, which suggests that a fucosidase must act before this galactosidase in the breakdown of the full plant N-glycan substrate. Analysis of the galactosidase activity against a soya-derived complex plant N-glycan structure confirmed that the Gal decorations could only be removed after the antennary fucoses have first been cleaved (Fig. 4*C*). This enzyme is also predicted to be periplasmic (Table 2).

Notably, this galactosidase did not have close homologues in other *Bacteroides* species and was highlighted in *B. massiliensis* by its association with B035DRAFT_00995^GH3^ xylosidase. Genes encoding putative β-1,3-galactosidases in other species with Group 2 PNGase enzymes were not obvious from the functional association analysis, suggesting the terminal galactose structures are likely targeted by a CAZyme unrelated to this GH2.

### Antennary fucose removal from plant N-glycans

Complex plant N-glycans are also often decorated with antennary α-1,4-fucose (Fig. 1*A*). The GH families that act to remove fucose in an exo-fashion include GH29, GH95, and GH151. GH29 enzymes typically act on α-1,3/4-linkages, but there are some examples of α-1,2-specific enzymes. Only a single GH29 was identified in *B. massiliensis* using functional association analysis (B035DRAFT_02132^GH29^) and this was shown to be specific for the core α-1,3 fucose. Therefore, to test the possibility of this species being able to degrade the antennary fucose structures we screened the activity of three further GH29 enzymes from *B. massiliensis*. All three displayed relatively broad activity against Lewis and fucosyllactose glycans (Fig. S11). In particular, all three were able to hydrolyse the α-1,4-fucose from Lewis A trisaccharide, which is the epitope found in plant complex N-glycans, whereas B035DRAFT_02132^GH29^ was unable to do this.

The GH29 fucosidases were then assessed against soya bean derived N-glycans (Fig. 4*C*). B035DRAFT_00014^GH29^ and B035DRAFT_03357^GH29^ were able to remove the antennary fucose from the complex N-glycan structures. Interestingly, B035DRAFT_00409^GH29^ was unable to do this despite being active against Lewis A trisaccharide. Furthermore, incubation of B035DRAFT_00996^GH2^ against the soya bean N-glycans in combination with either B035DRAFT_00014^GH29^ or B035DRAFT_03357^GH29^, showed removal of the terminal galactose and demonstrates that this galactosidase can remove galactose from plant N-glycan structures.

Although this screen of GH29 activities in *B. massiliensis* is not exhaustive, it does demonstrate that there are multiple enzymes present in the genome with the capability to access the antenna fucose from plant N-glycans.

### Accessing the activity of B035DRAFT_00997^sulfatase^

Sulfated N-glycans have been observed in a wide variety of organisms ranging from animals to viruses (Fig. S13)(23-26). These decorations can take the form of GalNAc-6S, GalNAc-4S, Gal-3S, Gal-6S, and Man-6S(36). To our knowledge, sulfation of plant N-glycans has not yet been observed, however, with N-glycan sulfation being so widespread throughout other organisms it would be surprising if it was not also present in some plants.

A putative sulfatase gene adjacent to the genes for the xylosidase and galactosidase (B035DRAFT_00997) was assessed for activity against a variety of sulfated monosaccharides and oligosaccharides (Fig. S13). No activity could be observed against the tested substrates. It is possible that this enzyme has specificity for a substrate not tested here, but we were not able to test the full spectrum of possibilities.

### Degradation of high-mannose N-glycan structures

The degradation of complex N-glycans in *B. thetaiotaomicron* has previously been described(12). This work showed three GH92 enzymes BT3990, BT3991, and BT3994 would hydrolyse the terminal α1,2-, α1,3-, and the first α1,6-mannose from high mannose N-glycans, respectively, to leave a Manα-1,6Manβ-1,4GlcNac trisaccharide. Homologues of these three enzymes were adjacent to the plant-N-glycan degrading genes in *B. helcogenes*, therefore it appears that this species has the genes required to degrade high-mannose and plant complex N-glycans in the same place in the genome. Homologues of these enzymes were also traced throughout the other species assessed in this study and found to be well-preserved throughout. Phylogenetic analysis of all the GH92 enzymes from the functional analysis was carried out and these clustered into five groups (Fig S14). Three of these are likely the GH92 enzymes acting high-mannose N-glycans, one group are likely all α1,3-mannosidases that can accommodate β1,2-xylose (homologues of B035DRAFT_03340^GH92^) and one remains uncharacterised.

## Discussion

This study characterises the pathway for the degradation of plant N-glycans by a prominent member of the gut microbiota. This set of enzymes was identified through functional association using putative PNGase enzymes as a starting point. This work demonstrates that it is possible in some cases to find enzymes with particular activities without using gene upregulation methods and only using what is already known about CAZyme families. This is a useful demonstration because for many substrates, like for the plant complex N-glycans described here, it is not possible to perform gene upregulation studies to identify the link between a substrate and set of genes.

Here we presented the characterisation of five enzymes against plant complex N-glycans and two of these include crystal structures. The specificity of these enzymes also indicated the order in which they act *in vivo* (Fig. 6). B035DRAFT_03341^PNGase^ removes the plant N-glycan from the protein and then monosaccharides are removed sequentially from the non-reducing ends. Firstly, an α1,4-fucosidase removes the fucose to allow B035DRAFT_00996^GH2^ to remove the terminal galactose. A number of different GH20 enzymes have been identified previously that can remove the GlcNAc at this stage and homologues of these enzymes are present in many of *Bacteroides* species(9). B035DRAFT_03340^GH92^ is then able to remove the α1,3-mannose, followed by B035DRAFT_00995^GH3^ removing the β1,2-xylose, and B035DRAFT_02132^GH29^ removing the core α1,3-fucose. This leaves a Manα-1,6Manβ-1,4GlcNacβ-1,4GlcNac tetrasaccharide.

In addition to providing understanding of how plant N-glycans are hydrolysed in the human gut, the experiments using insect glycoproteins indicate that this type of N-glycan may also be used as a nutrient source for the human gut microbiota. Insects have been a human food source for centuries for some populations, they are common in other primate diets, and there is an increased interest for this western culture largely for environmental and sustainability issues(27).

This report provides new methods to analyse plant complex N-glycans. It also provides more options for modifying proteins decorated with plant and insect N-glycans, such as biopharmaceuticals. One of the biggest potential uses of the CAZymes identified in this study would be in the production of pharmaceutical proteins in different plant species. Successful examples of this type of production include antibodies (“plantibodies”), collagen, vaccines, and enzymes, which can be produced in maize, rice, tobacco, flax, or strawberry(28). Monoclonal antibodies are potent treatments for a number of human diseases, including cancer and COVID-19(29). Variation in the composition of the N-glycans decorating the antibodies have been seen to affect the function of these biological therapeutics(30). Therefore, increasing the options around being able to modify N-glycans post-production will increase the success rate of different candidates in plants and insects. It will also provide opportunities to reduce the allergenicity of plant-produced proteins.

## Methods

### Sources of glycans and glycoproteins

Glycoproteins bovin α_1_acid glycoprotein, bovine fetuin, bovine RNaseB, horseradish peroxidase, bee venom phospholipase A_2_, and *p*-Nitrophenyl (pNP) monosaccharides were obtained from Sigma. Defined oligosaccharides were purchased from Carbosynth. Three Avastin batch samples were provided by Leaf Expression Systems and Sf9 insect cells were a gift from Professor Wyatt Yue and Dr Thomas McCorvie. The isolation of papaya and soya N-glycans is described in detail below.

### Bacterial strains

The *Bacteroides* strains used were: *B. fragilis* NCTC9342 and *B. massiliensis* DSM17679. *B. fragilis* was grown on tryptone-yeast-extract-glucose medium with the addition of haematin(31) and *B. massiliensis* was grown of chopped meat broth(32, 33) and both were inoculated from glycerol stocks. Genomic DNA was prepared using a 5 ml culture.

### Cloning, expression and purification of recombinant proteins

DNA encoding the appropriate genes (excluding the signal sequences) were amplified from genomic DNA using appropriate primers and cloned into pET28b (Novagen). Recombinant plasmids were transformed into TUNER (Novagen) cells in LB broth containing 10 μg/ml kanamycin at 37 °C shaking at 180 rpm. One litre cultures were grown to mid-exponential phase in 2 litre baffled flasks, cooled to 16 °C and isopropyl β-D-thiogalactopyranoside (IPTG) added to a final concentration of 0.2 mM. These cells were then incubated for 16 hours at 16 °C in an orbital shaker at 150 rpm. Recombinant His-tagged protein was purified from cell-free extracts using immobilised metal affinity chromatography (IMAC using Talon resin; Clontech) as described previously (34). The purity and size of the proteins were checked using SDS-PAGE and their concentrations determined using absorbance at 280 nm (NanoDrop 2000c; Thermo Scientific) and their molar extinction coefficients(35).

### Recombinant enzyme assays

The activities of the recombinant enzymes were typically assessed in 20 mM MOPS pH 7, at 37 °C, with a final glycoprotein concentration of 20 mg/ml and a final enzyme concentration of 1 μM. The bee venom phospholipase A_2_ assay was carried out at 0.5 mg/ml, with 4 μg loaded on a SDS-PAGE gel. The SDS-PAGE gel used weas a pre-cast 8-16 % gradient (Bio-Rad) and initially stained using Pro-Q™ Emerald 300 glycoprotein staining kit to highlight glycoproteins and subsequently stained with coomassie to visualise total protein. For overnight assays, defined oligosaccharides were incubated at a final concentration of 1 mM in the presence of 3 μM of enzyme.

### Thin-layer chromatography

For defined oligosaccharides, 3 μl of an assay containing 1 mM substrate was spotted on to silica plates. For assays against mucin, this was increased to 9 μl. The plates were resolved in running buffer containing butanol/acetic acid/water (2:1:1) and stained using a diphenylamine-aniline-phosphoric acid stain(36).

### Procainamide labeling

Procainamide labelling was performed by reductive amination using a procainamide labelling kit containing sodium cyanoborohydride as a reductant (Ludger). Excess reagents were removed with S cartridges (Ludger). Cartridges were conditioned successively with 1mL of DI water, 5 mL of 30 % acetic acid (v/v), and 1 mL of acetonitrile. Procainamide labelled samples were then spotted on the cartridge and allowed to adsorb for 15 min. The excess dye was washed with acetonitrile. Labelled N-glycans were eluted with 1 mL of DI water.

### Liquid chromatography-fluorescence detection-electrospray-mass spectrometry analysis of procainamide labelled glycans

Procainamide labelled glycans were analysed by LC-FLR-ESI-MS. Here, 25 μl of each sample (prepared in 24:76 water: acetonitrile solution) was injected into a Waters ACQUITY UPLC Glycan BEH Amide column (2.1 × 150 mm, 1.7 μm particle size, 130 Å pore size) at 40°C on a Dionex Ultimate 3000 UHPLC instrument with a fluorescence detector (λ_ex_ = 310 nm, λ_em_= 370 nm) attached to a Bruker Amazon Speed ETD. Mobile phase A was a 50 mM ammonium formate solution (pH 4.4) and mobile phase B was neat acetonitrile. Analyte separation was accomplished by gradients running at a flow rate of 0.4 ml/min from 85 to 57 % mobile phase B over 115 min and from 85 to 62 % over 95 min for mucin and keratan samples, respectively. The Amazon speed was operated in the positive sensitivity mode using the following settings: source temperature, 180 °C; gas flow. 41 min^-1^; capillary voltage, 4,500 V; ICC target, 200,000; maximum accumulation time, 50.00 ms; rolling average, 2; number of precursor ions selected, 3; scan mode, enhanced resolution; mass range scanned, 400 to 1,700.

### Analysis of mass spectrometry data

Mass spectrometry of procainamide-labelled glycans was analysed using Bruker Compass Data Analysis Software and GlycoWorkbench(37). Glycan compositions were elucidated on the basis of MS^2^ fragmentation and previously published data.

### Enzymatic papaya N-glycan release and 2-AB labeling

The applied N-glycan release method is based on the procedures described by Wilson et al. (5) and Du et al. (38). Briefly, papaya (ca. 2 g) were blended in a 2 ml glass homogenizer, transferred into a 2 ml centrifuge tube and then centrifuged (20000g for 20 min at 4°C). One mL of the clear supernatant was mixed with 1 mL of aqueous trichloroacetic acid (TCA) solution (2 M) and then centrifuged (20000g for 30 min at 4°C). The resulting pellet was once washed with 1 ml distilled water to remove TCA. The pellet was then re-suspended in 70 μl of distilled water, 28.5 μl of MES buffer (200 mM, pH 7.0) and 12.5 μl of denaturation solution (2% SDS (w/V) and 2-mercaptoethanol (1 M) in water) were added. The mixture was heated at 100°C for 5 min in a heating block. After cooling, 19 μl of a Triton solution (10% w/V) was added followed by the addition of 100 μlL of purified PNGase, and the mixture was incubated overnight at 37°C. The supernatant was collected by centrifugation at 12000 rpm for 20 min and purified with Supelclean ENVI Carb solid-phase extraction (SPE, 500 mg bed volume) columns. The SPE columns was activated with 3 ml of 80% acetonitrile containing 0.1% trifluoroacetic acid (TFA, V/V) and equilibrated with the same volume of distilled water. The sample was loaded to the SPE column and then washed with 3 mL of distilled water. N-glycans were eluted with 20% and 40% acetonitrile containing 0.1% TFA (v/v), collected, dried, and labeled with 2-AB. To do so, an aliquot (10 μl) of 2-AB labelling solution (35 mM of 2-AB and 0.1 M of sodium cyanoborohydride in dimethyl sulfoxide/acetic acid (7:3 v/v)) was added, and the mixture was incubated at 65°C for 4 h.

### Papaya N-glycan separation

Chromatographic separation of oligosaccharides was carried using a Nexera UPLC system (Shimadzu Corporation, Kyoto, Japan), consisting of a DGU-20A5R degasser unit, a LC-30AD pump, a SIL-30AC autosampler, and a RF-20Axs fluorescence detector (set at 330 nm excitation and 420 nm emission) adapted from Guo *et* al. (39). Briefly, the analyses were performed using an Acquity BEH Glycan column (Waters 1.7 μm, 2.1 × 150 mm). The mobile phases consisted of NH_4_COOH (pH 4.5, 50 mM) in water and acetonitrile for solvents A and B, respectively. The elution methods were set as follows: a linear gradient of 95-78 % B was applied from 0-6 min at a flow rate of 0.5 ml/min; 78-70 % B from 6-20 min at 0.5 ml/min; 70-0 % B in 1 min at 0.25 ml/min; held for 2 min at 0.25 ml/min; 0-95 % B in 2 min at 0.25 ml/min; held for 1.5 min at 0.25 ml/min; the flow rate was then increased from 0.25 to 0.5 mL/min from 26.5-29.5 min; and finally the column was equilibrated with 95 % B for 3.5 min at 0.5 mL/min before the next sample was injected.

### MALDI-TOF-MS analysis of papaya samples

UPLC fractions corresponding to selected fluorescence peaks were collected in 2 ml centrifuge tubes, dried using vacuum centrifugation (Speedvac), and re-dissolved in 10 μl of distilled water. 1 μl fractions of this samples were then analysed using a Bruker Auto-flex Speed (Bruker Daltonics, Bremen, Germany) MALDI-TOF-MS spectrometer (equipped with a 1000 Hz Smartbeam laser). Samples were overlaid with 1 μl of 2,5-dihydroxybenzoic acid (DHB) matrix (10 mg/ml DHB in 70% (v/v) aqueous acetonitrile solution). Mass spectra were analysed using the Bruker FlexAnalysis software version 3.3.80, and N-glycan masses were calculated using the GlycoWorkbench software tool (37).

### N-Glycan release from soya protein and MALDI analysis

Analytical release of N-glycans from soy proteins was conducted as follows. 1 g of soy protein isolate (purchased from a local supermarket) was washed with deionized (DI) water (3x 10 mL), with centrifugation (10 min X 2500 g) in between each wash. The resulting pellet was homogenized with 20 mL of DI water to form a slurry. 100 μL of the slurry was dried down by vacuum centrifugation and resuspended in 25 μl of 50 mM NaH_2_PO_4_-Na_2_HPO_4_ buffer pH 7.5 and boiled for 5 min. Control samples were digested with PNGase F (1 μl, 5 mU, QA-Bio) For Pngase-003341 the final enzyme concentration was 1 μM. Samples were incubated for 12 h at 37 °C. 100 μl of DI water was added to dilute the sample before MALDI-MS analysis using a Bruker Auto-flex Speed (Bruker Daltonics, Bremen, Germany). The spectrometer was operated in positive ion mode. Spectra were acquired in the mass range 900–3500 m/z at a laser intensity of 50%. The Mass Spectrometry (MS) data were further processed using Flex analysis3.5; sample preparation was as follows 0.5 μl of Super-DHB matrix (50 mg/mL in (50:50 [v/v] H_2_O: acetonitrile)), was spotted on a ground steel target, 0.5 μl the sample was added on top and allowed to dry. 20 μl of aliquots of the diluted samples were dried down by vacuum centrifugation and labelled with procainamide prior to UHPLC-MS analysis.

### Isolation of plant complex N-glycans from soya proteins

A total of 20 g of soya protein isolate were processed as follows: 2.5 g of soy protein isolate were placed in 50 mL centrifuge tubes, for a total of eight tubes. To each tube 25 mL,50 mM NaH_2_PO_4_-Na_2_HPO_4_ pH 6.0, 0.05 % NaN_3_ buffer was added (1:10 solid-liquid ratio), and the samples were denatured by boiling for 5 min at 100 °C. The tubes were allowed to cool down and 60 μL of PNGaseL (2 mg/mL) were added and allowed to incubate for 2d at 37 °C. After release samples were centrifuged (30 min 2500 g). and the pellet was washed thrice with DI water (20 mL), supernatants were combined and concentrated by rotary evaporation to 20 mL. Acetone was added to achieve a concentration of 50% (v/v) and allowed to cool at –20 °C for 1h to precipitate proteins, the supernatant was separated by centrifugation (30 min X 2500 g). The pellet was washed twice with 10 mL of 50% acetone, and the washings were combined with the supernatant and concentrated to dryness using a rotary evaporator. The dry residue was resuspended in 5 mL of H_2_O +0.1 % TFA (v/v) and loaded to a 5 g C18 Supelclean™ LC18-SPE cartridge pre-conditioned with 50 mL methanol, followed by 50 mL H_2_O + 0.1 % (v/v) TFA to remove residual proteins and other hydrophobic contaminants and washed with 50 mL of H_2_O +0.1% TFA, in 5 mL of DI water and loaded to a 2x High prep 26/10 Sephadex-G25 columns. Glycan containing fractions were detected with MALDI-MS and dried down using a SpeedVac vacuum concentrator. Dried glycan samples were resuspended in 70% acetonitrile. Injections of 1.5 mL were applied to a semi-preparative HILIC-column (TSKgel-amide-80, 7.8 i. d x300 mm, 10 μm, Tosoh Biosciences) at 50°C on a Dionex Ultimate 3000 UHPLC with an automatic fraction collector. Elution was performed at a flow rate of 2.0 mL/min. Solvent A was 50 mM ammonium formate (pH 4.4), solvent B was acetonitrile. The column was equilibrated with 70 % solvent B. The gradient elution parameters were 60-48 % solvent B with a linear gradient over 68 min. Detection was carried out at 214 nm and glycan peaks were collected as they eluted from the column, MALDI-MS was used for fraction identification. The fractions containing plant complex N-glycans were combined and concentrated to dryness, salts were removed using a 2x High prep 26/10 Sephadex-G-25 column and subjected to an additional round of purification through the TSKgel-amide-80 column. The fractions containing plant complex N-glycans were combined once again and salts were removed using 2x High prep 26/10 Sephadex-G-25 column.

### Exoglycosidase digestion of plant complex glycans from soya proteins

150 pmol of the purified soya protein was labelled with procainamide and used as a substrate for exoglycosidase reactions. For each exoglycosidase digestion, 10 pmol of procainamide labelled soya protein were used. Digestions were carried out in 50 mM sodium phosphate buffer, pH 7 in a final volume of 10 μl for 18h at 37°C. Exoglycosidases were used at a final concentration of 1 μM. After incubation, glycans were purified using a Ludger LC-EXO-96 plate and eluted in 200 μl of water. The samples were then dried down by vacuum centrifugation, solubilised in 30 μl of water and analysed by UHPLC-MS.

### Crystallization

B035DRAFT_03341^PNGase^ and B035DRAFT_03340^GH92^ were initially screened using commercial kits (Molecular Dimensions, Qiagen and Hampton Research). The protein concentrations were 20 mg/ml. The drops, composed of 0.1 μl or 0.2 μl of protein solution plus 0.1 μl of reservoir solution, were set up a Mosquito crystallization robot (SPT Labtech). The sitting drop method was used and the plates were incubated at 20 °C. The crystallisation condition for B035DRAFT_03340^PNGase^ was condition B12 in JCSG screen part I (Qiagen). B035DRAFT_03341^GH92^ was condition E12 in Index screen (Hampton Research). The samples were cryoprotected with the addition of 20 % PEG 400 to the crystallisation condition.

### Data collection, structure solution, model building, refinement and validation

Diffraction data were collected at the synchrotron beamlines I03 and I04 of Diamond light source (Didcot, UK) at a temperature of 100 K. The data set was integrated with dials(40) or XDS(41) via XIA(42) and scaled with Aimless(43). The space group was confirmed with Pointless. The phase problem was solved by molecular replacement with Phaser(44) using PDB file 2WVX and 4R4Z as search models for B035DRAFT_03340^PNGase^ and B035DRAFT_03341^GH92^ respectively. While the initial solution Rfactors were very poor (over 50%) for B035DRAFT_03340^PNGase^ the electron density map was interpretable. The automated model building program task CCP4build on CCP4cloud(45) delivered a model with Rfactors below 20 %. The models were refined with refmac(46) and manual model building with COOT(47). The final model was validated with COOT(47) and MolProbity(48). Other program used were from the CCP4 suite(45). Processing and refinement statistics are reported in table XX. Models and data were deposited to the Protein Data Bank with the codes 7ZGN and 7ZGM for B035DRAFT_03340^PNGase^ and B035DRAFT_03341^GH92^, respectively. Figures were made with PyMol(49).

### Bioinformatics

Putative signal sequences were identified using SignalP 5.0(50). Sequence identities were determined using Clustal Omega using full sequences(51). The IMG database was used to analyse synteny between different species(52). The CAZy database (www.cazy.org) was used as the main reference for CAZymes(53). Alignments and phylogenetic trees were completed in SeaView(54). To determine the boundaries between different modules in a protein Pfam(55) and SMART(56, 57) were used.

## Acknowledgements

We thank Carl Morland (Newcastle University, UK) for his expert technical assistance. The insect cells were a kind gift from Professor Wyatt Yue and Dr Thomas McCorvie. We would like to thank Diamond Light Source (Oxfordshire, UK) for beamtime (proposal mx13587) and staff of beamline I03 and I04. The work was funded by BBSRC/Innovate UK IB catalyst award ‘Glycoenzymes for Bioindustries’ (BB/M029018/1) and National Natural Science Foundation of China (grant numbers 31471703, A0201300537 and 31671854 to J.V. and L.L.).

## Author contributions

L.I.C. and D.N.B designed the research. L.I.C predominantly characterised the enzymes and set up crystal trays. P.A.U. gathered most of the LC-FLD-ESI-MS data. A.B. harvested the crystals, gathered the data, and solved the crystal structures. For the soya protein work: P.A.U., J.M.MD., and D.I.R.S. isolated the soya protein N-glycans, performed enzymes assays against this substrate, and gathered the LC-FLD-ESI-MS data. S.T.B gathered data against insect-derived glycoproteins. For the papaya work: Z.P.C. performed the experiments and gathered the data. Z.P.C., L.L., and J.V. analysed parts of the data. L.L. and J.V. coordinated parts of the project. L.I.C. and D.N.B analysed the data and wrote the paper with everyone contributing to the methods section.

## Data availability

The crystal structures and data are deposited in the Protein Data Bank under the accession numbers 7ZGM and 7ZGN. The other data that supports the findings in this paper are available upon request from the corresponding authors.

**Supplementary Table S1.**
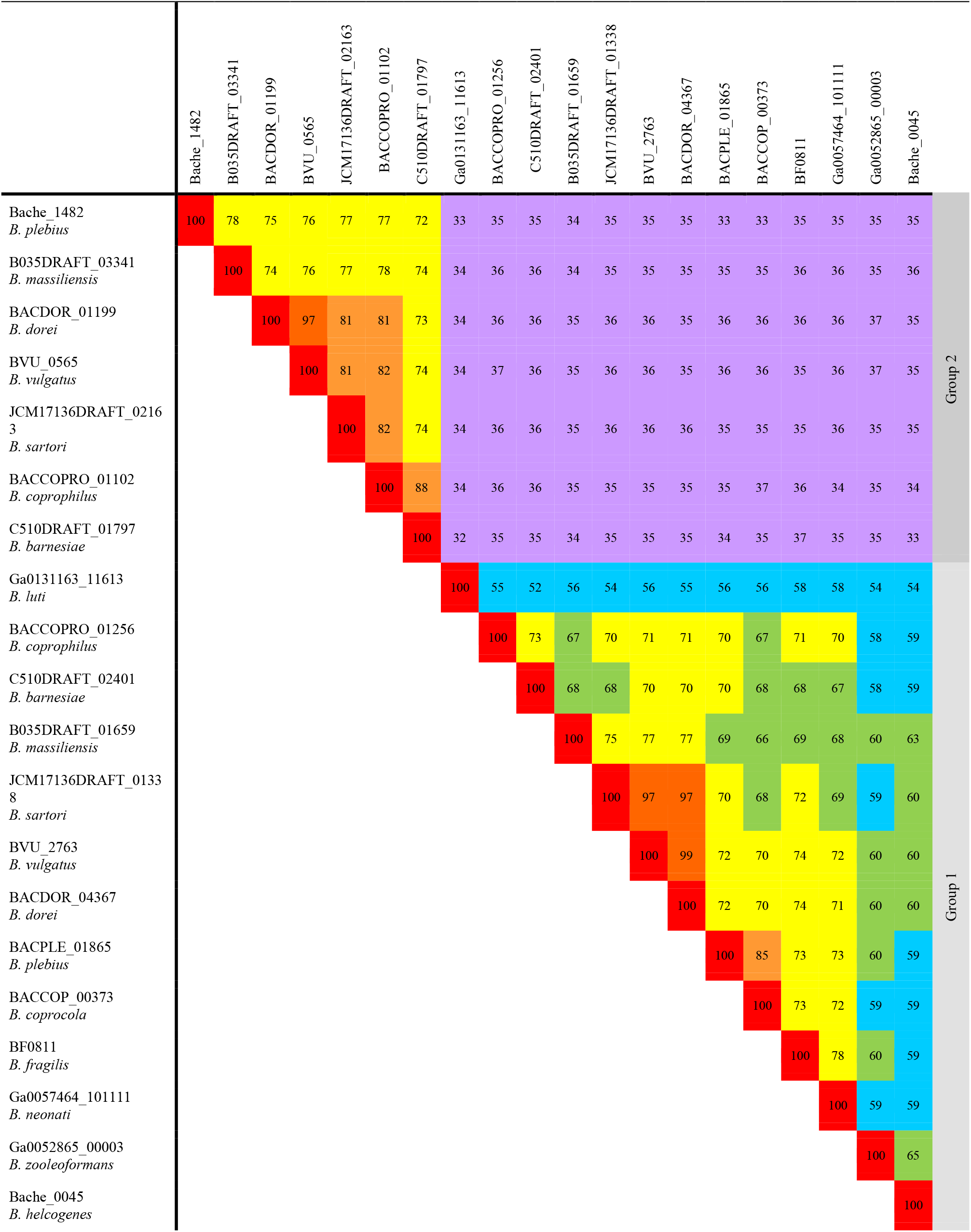
Percentage identity between putative PNGase enzymes from species of *Bacteroides*.

|  | Bache_1482 | B035DRAFT_03341 | BACDOR_01199 | BVU_0565 | JCM17136DRAFT_02163 | BACCOPRO_01102 | C510DRAFT_01797 | Ga0131163_11613 | BACCOPRO_01256 | C510DRAFT_02401 | B035DRAFT_01659 | JCM17136DRAFT_01338 | BVU_2763 | BACDOR_04367 | BACPLE_01865 | BACCOP_00373 | BF0811 | Ga0057464_101111 | Ga0052865_00003 | Bache_0045 |  |
| --- | --- | --- | --- | --- | --- | --- | --- | --- | --- | --- | --- | --- | --- | --- | --- | --- | --- | --- | --- | --- | --- |
| Bache_1482<br>*B. plebius* | 100 | 78 | 75 | 76 | 77 | 77 | 72 | 33 | 35 | 35 | 34 | 35 | 35 | 35 | 33 | 33 | 35 | 35 | 35 | 35 | Group 2 |
| B035DRAFT_03341<br>*B. massiliensis* |  | 100 | 74 | 76 | 77 | 78 | 74 | 34 | 36 | 36 | 34 | 35 | 35 | 35 | 35 | 35 | 36 | 36 | 35 | 36 |  |
| BACDOR_01199<br>*B. dorei* |  |  | 100 | 97 | 81 | 81 | 73 | 34 | 36 | 36 | 35 | 36 | 36 | 35 | 36 | 36 | 36 | 36 | 37 | 35 |  |
| BVU_0565<br>*B. vulgatus* |  |  |  | 100 | 81 | 82 | 74 | 34 | 37 | 36 | 35 | 36 | 36 | 35 | 36 | 36 | 35 | 36 | 37 | 35 |  |
| JCM17136DRAFT_02163<br>*B. sartori* |  |  |  |  | 100 | 82 | 74 | 34 | 36 | 36 | 35 | 36 | 36 | 36 | 35 | 35 | 35 | 36 | 35 | 35 |  |
| BACCOPRO_01102<br>*B. coprophilus* |  |  |  |  |  | 100 | 88 | 34 | 36 | 36 | 35 | 35 | 35 | 35 | 35 | 37 | 36 | 34 | 35 | 34 |  |
| C510DRAFT_01797<br>*B. barnesiae* |  |  |  |  |  |  | 100 | 32 | 35 | 35 | 34 | 35 | 35 | 35 | 34 | 35 | 37 | 35 | 35 | 33 |  |
| Ga0131163_11613<br>*B. luti* |  |  |  |  |  |  |  | 100 | 55 | 52 | 56 | 54 | 56 | 55 | 56 | 56 | 58 | 58 | 54 | 54 | Group 1 |
| BACCOPRO_01256<br>*B. coprophilus* |  |  |  |  |  |  |  |  | 100 | 73 | 67 | 70 | 71 | 71 | 70 | 67 | 71 | 70 | 58 | 59 |  |
| C510DRAFT_02401<br>*B. barnesiae* |  |  |  |  |  |  |  |  |  | 100 | 68 | 68 | 70 | 70 | 70 | 68 | 68 | 67 | 58 | 59 |  |
| B035DRAFT_01659<br>*B. massiliensis* |  |  |  |  |  |  |  |  |  |  | 100 | 75 | 77 | 77 | 69 | 66 | 69 | 68 | 60 | 63 |  |
| JCM17136DRAFT_01338<br>*B. sartori* |  |  |  |  |  |  |  |  |  |  |  | 100 | 97 | 97 | 70 | 68 | 72 | 69 | 59 | 60 |  |
| BVU_2763<br>*B. vulgatus* |  |  |  |  |  |  |  |  |  |  |  |  | 100 | 99 | 72 | 70 | 74 | 72 | 60 | 60 |  |
| BACDOR_04367<br>*B. dorei* |  |  |  |  |  |  |  |  |  |  |  |  |  | 100 | 72 | 70 | 74 | 71 | 60 | 60 |  |
| BACPLE_01865<br>*B. plebius* |  |  |  |  |  |  |  |  |  |  |  |  |  |  | 100 | 85 | 73 | 73 | 60 | 59 |  |
| BACCOP_00373<br>*B. coprocola* |  |  |  |  |  |  |  |  |  |  |  |  |  |  |  | 100 | 73 | 72 | 59 | 59 |  |
| BF0811<br>*B. fragilis* |  |  |  |  |  |  |  |  |  |  |  |  |  |  |  |  | 100 | 78 | 60 | 59 |  |
| Ga0057464_101111<br>*B. neonati* |  |  |  |  |  |  |  |  |  |  |  |  |  |  |  |  |  | 100 | 59 | 59 |  |
| Ga0052865_00003<br>*B. zooleoformans* |  |  |  |  |  |  |  |  |  |  |  |  |  |  |  |  |  |  | 100 | 65 |  |
| Bache_0045<br>*B. helcogenes* |  |  |  |  |  |  |  |  |  |  |  |  |  |  |  |  |  |  |  | 100 |  |

**Supplementary Table S2.** Signal peptide predictions of the enzymes characterised in this study.

| Species | Locus Tag | Enzyme family | Predicted signal peptide (using SigP5.0) |
| --- | --- | --- | --- |
| *B. massiliensis* | B035DRAFT_03341 | PNGase | SPI |
| *B. fragilis* | BF0811 | PNGase | SPI |
| *B. massiliensis* | B035DRAFT_03340 | GH92 | SPII |
| *B. massiliensis* | B035DRAFT_00995 | GH3 | SPI |
| *B. massiliensis* | B035DRAFT_00996 | GH2 | SPI |
| *B. massiliensis* | B035DRAFT_00997 | Sulfatase | SPI |
| *B. massiliensis* | B035DRAFT_02132 | GH29 | SPI |
| *B. massiliensis* | B035DRAFT_00014 | GH29 | None |
| *B. massiliensis* | B035DRAFT_00409 | GH29 | SPI |
| *B. massiliensis* | B035DRAFT_03357 | GH29 | SPI |

**Supplementary Table S3.** Data statistics and refinement details.

| Data statistics* |  |  |
| --- | --- | --- |
|  | B035DRAFT_03340 | B035DRAFT_03341 |
| Beamline | I04 | I03 |
| Date | 20/01/17 | 22/05/17 |
| Wavelength (Å) | 0.979 | 0.979 |
| Resolution (Å) | 66.29 – 1.43 (1.45 – 1.43) | 45.09 – 1.95 (1.99 – 1.95) |
| Space group | P2<sub>1</sub>2<sub>1</sub>2<sub>1</sub> | P2<sub>1</sub>2<sub>1</sub>2<sub>1</sub> |
| Unit-cell parameters |  |  |
| a (Å) | 65.94 | 59.05 |
| b (Å) | 84.35 | 100.55 |
| c (Å) | 132.61 | 180.37 |
| $\alpha = \beta = \gamma$ (°) | 90.00 | 90.00 |
| Unit-cell volume (Å<sup>3</sup>) | 737293 | 1070942 |
| Solvent content (%) | 47 | 43 |
| No. of measured reflections | 959206 (38443) | 573096 (31978) |
| No. of independent reflections | 135130 (6477) | 79080 (4454) |
| Completeness (%) | 98.8 (97.1) | 99.9 (99.6) |
| Redundancy | 7.1 (5.9) | 7.2 (7.2) |
| CC<sub>1/2</sub> (%) | 0.998 (0.502) | 0.999 (0.517) |
| $\langle I \rangle / \langle \sigma(I) \rangle$ | 9.7 (1.2) | 12.6 (1.7) |
| Refinement statistics* |  |  |
| Rwork (%) | 13.32 | 18.92 |
| Rfree<sup>#</sup> (%) | 17.22 | 23.95 |
| No. of non-H atoms |  |  |
| No. of protein, atoms | 5661 | 4321 |
| No. of solvent atoms | 646 | 336 |
| No. of ion atoms | 17 | 0 |
| R.m.s. deviation from ideal values |  |  |
| Bond angle (°) | 1.22 | 1.77 |
| Bond length (Å) | 0.011 | 0.011 |
| Average B factor (Å<sup>2</sup>) |  |  |
| Protein | 20 | 47 |
| Solvent | 33 | 42 |
| Ions | 26 | NA |
| Ramachandran plot<sup>+</sup>, residues in |  |  |
| Most favoured regions (%) | 97.01 | 96.1 |
| PDB code |  |  |

\*(Values in parenthesis are for the highest resolution shell).
<sup>#</sup>5% of the randomly selected reflections excluded from refinement.
<sup>+</sup>Calculated using MOLPROBITY.

**Supplementary Table S4.** Percentage identity between putative plant N-glycan specific α-1,3-mannosidase GH92 enzymes.

|  | Ga0057464_101116 | BACCOPRO_02654 | C510DRAFT_01523 | Bache_1481 | B035DRAFT_03340 | BVU_0555 | JCM17136DRAFT_01210 | BACDOR_01210 |
| --- | --- | --- | --- | --- | --- | --- | --- | --- |
| Ga0057464_101116<br>*B. neonati* | 100 | 73 | 70 | 71 | 72 | 72 | 72 | 72 |
| BACCOPRO_02654<br>*B. coprophilus* |  | 100 | 83 | 78 | 80 | 80 | 80 | 80 |
| C510DRAFT_01523<br>*B. barnesiae* |  |  | 100 | 77 | 78 | 79 | 78 | 78 |
| Bache_1481<br>*B. helcogenes* |  |  |  | 100 | 82 | 83 | 82 | 82 |
| B035DRAFT_03340<br>*B. massiliensis* |  |  |  |  | 100 | 87 | 88 | 88 |
| BVU_0555<br>*B. vulgatus* |  |  |  |  |  | 100 | 94 | 95 |
| JCM17136DRAFT_01210<br>*B. sartori* |  |  |  |  |  |  | 100 | 96 |
| BACDOR_01210<br>*B. dorei* |  |  |  |  |  |  |  | 100 |

**Supplementary Table S5.** Percentage identity between putative plant N-glycan specific β-1,2-xylosidase GH3 enzymes.

|  | C510DRAFT_01523 | BACCOPRO_02654 | Ga0131163_10288 | B035DRAFT_03340 | Bache_1482 | BVU_0555 |
| --- | --- | --- | --- | --- | --- | --- |
| C510DRAFT_01523<br>*B. barnesiae* | 100 | 80 | 61 | 65 | 67 | 67 |
| BACCOPRO_02654<br>*B. coprophilus* |  | 100 | 60 | 64 | 66 | 65 |
| Ga0131163_10288<br>*B. luti* |  |  | 100 | 66 | 63 | 62 |
| B035DRAFT_03340<br>*B. massiliensis* |  |  |  | 100 | 76 | 75 |
| Bache_1482<br>*B. plebius* |  |  |  |  | 100 | 84 |
| BVU_0555<br>*B. vulgatus* |  |  |  |  |  | 100 |

**Supplementary Table S6.** Percentage identity between putative plant N-glycan specific α-1,3-fucosidase GH29 enzymes.

|  | C510DRAFT_00332 | BACCOPRO_02242 | JCM17136DRAFT_003055 | Bache_1479 | B035DRAFT_02132 | Bache_1477 | C510DRAFT_00333 |
| --- | --- | --- | --- | --- | --- | --- | --- |
| C510DRAFT_00332<br>*B. barnesiae* | 100 | 80 | 74 | 77 | 77 | 69 | 67 |
| BACCOPRO_02242<br>*B. coprophilus* |  | 100 | 73 | 74 | 75 | 67 | 67 |
| JCM17136DRAFT_003055<br>*B. sartori* |  |  | 100 | 78 | 79 | 67 | 65 |
| Bache_1479<br>*B. helcogenes* |  |  |  | 100 | 80 | 70 | 67 |
| B035DRAFT_02132<br>*B. massiliensis* |  |  |  |  | 100 | 71 | 66 |
| Bache_1477<br>*B. helcogenes* |  |  |  |  |  | 100 | 69 |
| C510DRAFT_00333<br>*B. barnesiae* |  |  |  |  |  |  | 100 |

