## Supplemental Figures for "Plant N-glycan breakdown by human gut *Bacteroides*"

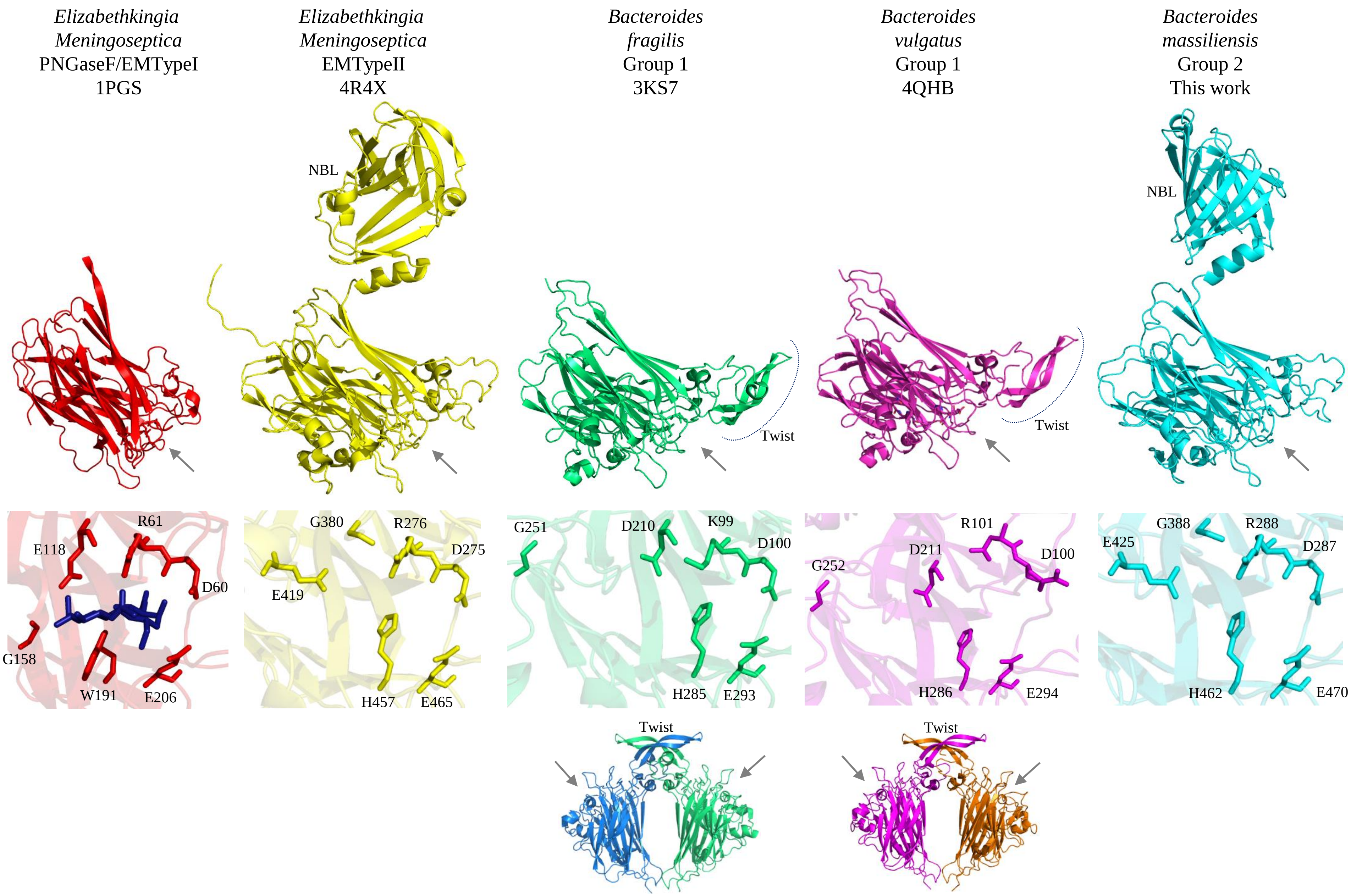


**Supplemental Figure S1. Crystal structures of PNGase enzymes from *Elizabethkingia meningoseptica* and *Bacteroides* species.** The catalytic domains are lined up and placed in the same orientation for comparison and the grey arrow shows the location of the active site.. The additional N-terminal bowl-like domain from EMTypeII and B035DRAFT_03341$^{PNGase}$ are labelled NBL. The key active site residues from the different enzymes are also shown as sticks. The PNGases from *B. fragilis* and *B. vulgatus* have residues D210 and D211, respectively, that are equivalent to E118 in PNGaseF/EMTypeI. Conversely, B035DRAFT_03341$^{PNGase}$ has G388 and E425, which are equivalent to G380 and E419 in EMTypeII, and form a pocket the likely allows the accommodation of the core α-1,3-fucose found in plant N-glycans. The small β-sheet twist present in the *B. fragilis* and *B. vulgatus* structures is labelled and the way two of these interact is shown.

```
EmenI        ------------------------------------------------------------ 0
Blut11613    ------------------------------------------------------------ 0
Bcopp01256   ------------------------------------------------------------ 0
Bbar02401    ------------------------------------------------------------ 0
Bmas01659    ------------------------------------------------------------ 0
Bsar01338    ------------------------------------------------------------ 0
Bvul2763     ------------------------------------------------------------ 0
Bdor04367    ------------------------------------------------------------ 0
Bple01865    ------------------------------------------------------------ 0
Bcopc00373   ------------------------------------------------------------ 0
Bfrag0811    ------------------------------------------------------------ 0
Bneo101111   ------------------------------------------------------------ 0
Bzoo00003    ------------------------------------------------------------ 0
Bhel0045     ------------------------------------------------------------ 0
EmenII       MLFFLPLLKTNLM--QKILLCSLIT---------GAQMIFAQTYEITYQNSFEGKINPNQ 49
Bhel1482     ------------------MLCILTLSTTISAQNMQKKLKNAKGIEVIYRSVYKGKTIPGQ 42
Bmas03341    -------MRRTGIRNAWLLMLCLLTVSAAAAQNLQKKVKNAKGIEVIYQSSYKGKIRPGQ 53
Bdor01199    -------MKRTILKDVGIGLCFLLSTGTIYAQNYPRKVKNAQGIEVTYQSNYKGRVRPGH 53
Bvul0565     -------MKRTILKNVGIGLCFLLSTGTVCAQNYPGKVKNAQGIEVTYQSNYKGKARPGH 53
Bsar02163    -------MKNFTMKELWLLLCCLMAVSAIHAQNYSKKVKNAKGIEVTYQSSYKGKVRPGY 53
Bcopp01102   ------------------------------------------------------------ 0
Bbar01797    -------MKHLNVKSCCLMVCCLLTVTTAEAENIRKKVKNAPGIEVTYQSSYKGKVAPGQ 53
```

Group 1

Group 2

N-terminal bowl-like domain

```
EmenI        ------------------------------------------------------------ 0
Blut11613    ------------------------------------------------------------ 0
Bcopp01256   ------------------------------------------------------------ 0
Bbar02401    ------------------------------------------------------------ 0
Bmas01659    ------------------------------------------------------------ 0
Bsar01338    ------------------------------------------------------------ 0
Bvul2763     ------------------------------------------------------------ 0
Bdor04367    ------------------------------------------------------------ 0
Bple01865    ------------------------------------------------------------ 0
Bcopc00373   ------------------------------------------------------------ 0
Bfrag0811    ------------------------------------------------------------ 0
Bneo101111   ------------------------------------------------------------ 0
Bzoo00003    ------------------------------------------------------------ 0
Bhel0045     ------------------------------------------------------------ 0
EmenII       NHIISITNSDKTLLFNEKIKNKK-----------ADFPFEVNEINRKNNEVSQFAFLNNN 98
Bhel1482     MQMTVCMD--QVALKNVLPPQEQSPETVGEPTPEIETPVTSNYIDYSSCQAYRLAKLPNG 100
Bmas03341    IKMTVSGN--QVALESVSPKGEK--ETATEGIREDKQPVIKNYIDYAGREAYKWAELPDG 109
Bdor01199    LLMTVSGD--RVSLTNVWSEQND----RPDPRPEDKTPVTGSYIDYTTRQAYRRAELPNG 107
Bvul0565     LLMTVSGD--RVSLTNVWPEQND----RPNPRPEDKTPVTGSYIDYTTRQAYRRAELPNG 107
Bsar02163    LLMTVSTD--RVSLENKRAESKQ--PNDNQVRPEDKTPVTGSYIDYTTCQSYRRAELPNG 109
Bcopp01102   ------------------------------------------------------------ 0
Bbar01797    VLMKVIGD--EVILTSLKPEGKP----AEEIREQDKAPVITNYIDYEACKSYKRAELPDG 107
```

Group 1

Group 2

```
EmenI        ------------------------------------------------------------ 0
Blut11613    ------------------------------------------------------------ 0
Bcopp01256   ------------------------------------------------------------ 0
Bbar02401    ------------------------------------------------------------ 0
Bmas01659    ------------------------------------------------------------ 0
Bsar01338    ------------------------------------------------------------ 0
Bvul2763     ------------------------------------------------------------ 0
Bdor04367    ------------------------------------------------------------ 0
Bple01865    ------------------------------------------------------------ 0
Bcopc00373   ------------------------------------------------------------ 0
Bfrag0811    ------------------------------------------------------------ 0
Bneo101111   ------------------------------------------------------------ 0
Bzoo00003    ------------------------------------------------------------ 0
Bhel0045     ------------------------------------------------------------ 0
EmenII       EIVKTSDNTILAKQEFKPTSETGKILGYNVKKAVTSVNSNTIEVWYTNDLKVKGGPS-IL 157
Bhel1482     KVISAATP-FRIGAGFT-EAGEGKHLGLNCKILRTSLRSNTIEVWYTNDIPFRGTPQANV 158
Bmas03341    KIISAATP-FEFGKGFT-PAGEGKHLGLNCKIARTSINSNTIEVWYTHDIPFRGTPQANV 167
Bdor01199    QVISAVTP-FEFGKGFT-QTGEGKHLGMNCKILRTSINSNTIEVWYTNDIPFRGTPQANV 165
Bvul0565     QVISAVTP-FEFGKGFT-QTGEGKHLGMNCKILRTSINSNTIEVWYTNDIPFRGTPQANV 165
Bsar02163    KIISAMTP-FELGRGFT-EKGEGKHLGLNCKIVRTSINSNTIEVWYTNEIPFRGTPQPNV 167
Bcopp01102   ------------------------------------------------------------ 0
Bbar01797    RIISAATP-FEYGKGFK-EVGTDKVLGLDCKILQTIINSNTIQVWYTTDIPFRGTPQANV 165
```

Group 1

Group 2

```
EmenI          ------------------------------------------------------------ 0
Blut11613      ------------------------------MNKNSFLSCGNAFIAMLFLVLAVCGQKLSA 30
Bcopp01256     --------------------------------MWKKVLP---YVAAGVAALSFAACGPKE 25
Bbar02401      --------------------------------MLKKILF---LAAACLFTWTGSAVEHKE 25
Bmas01659      ---------------------------MRRNNLLTFIAS---LALVTSFIMPANAAKHKE 30
Bsar01338      ------------------------------MNLLTFITP---LAVAATFMMPADAANHKE 27
Bvul2763       ------------------------------MNLLTFIAP---LAVAATFMMPADAANHKE 27   Group 1
Bdor04367      ------------------------------MNLLTFIAP---LAVAATFMMPADAANHKE 27
Bple01865      ----------------------------------MKIVS---MFVSLFVALNVCAAGHKE 23
Bcopc00373     ---------------MIYKNSNFINLKALEFNIDMRTIF---ILFSLLFTLTVNAASHKE 42
Bfrag0811      --------------------------------MNIRLTS---LFVSLFLSVPVWAGGHKN 25
Bneo101111     --------------------------------MNKNLYI---LLIALLSASMAWAGKHKE 25
Bzoo00003      ---------------------------------MNKITC---LLLSLFLLSSALSVSAKK 24
Bhel0045       ------------------------------MLMENKIL-----YLFVVLLGIPGLISARK 25
EmenII         GQDLGLVLKTVRNGSSVVEATSVKKIKALDDQS---------LFNGKNITEKDALTYKDM 208
Bhel1482       GVPDGLVLKVIRNGDMVQEASSISPLKKA-EN----------LLPTTWGEALDADDYQYT 207
Bmas03341      GVPDGLVLKVVRNGDMIQEASAITPLKKA-QA----------LLPDSWGEKMDAADYQYT 216
Bdor01199      GVPDGLVLRVVRNGDMIQEATHITPLKK-GKD----------VLPQSWGKSMDAADYQYT 214   Group 2
Bvul0565       GVPDGLVLRVVRNGDMIQEATHITPLKK-GKD----------VLPQSWGESMDAADYQYT 214
Bsar02163      GIPDGLVLKIVRNGDMVQEAILITPLKKE-AD----------LLPADWGEAMDAADYQYT 216
Bcopp01102     ---------------------------------------------------MDASDFQYT 9
Bbar01797      GVPDGLVLKVVRNGDTVQEAISINPEKETGTS----------LLPTSWGNVMDNADYQYT 215

EmenI          APADNTVNIKTFDKVKNAFGDGLSQS------AEGTFTFPADVTAVKTIK---------M 45
Blut11613      SNHNTEFVIKPFNETSVCFNSNDYPDK--VWEGDGLIRLDHGRIVIKKIRVPKFKQNVVV 88
Bcopp01256     YPAQGDCNLTVFQQERVRFCPDSIAN-YTAPDSNGVMRLVNGRILLKKITLPKYQRNIDV 84
Bbar02401      LPAKGDCSLQVFKQERVRFCPDSIGN-ITAPDADGVMRLVNGRILLKKIKLPEYQRNIKV 84
Bmas01659      LPALGDTQIQIFDKTNICFRPDSFAN-YTPASADGVIRLVNGRIILKKISLPDYKRNVRV 89
Bsar01338      LPALGNTHIQVFDKTPVCFRPDSFPN-YTPANADGVIRLVNGRIILKKITLPDYKRDVDV 86
Bvul2763       LPALGNTHIQVFDKTPVCFRPDSFPN-YTPANADGVIRLVNGRIILKKITLPDYKRDVDV 86   Group 1
Bdor04367      LPALGNTHIQVFDKTPVCFRPDSFPN-YTPANADGVIRLVNGRIILKKITLPDYKRDVDV 86
Bple01865      LPAKGDLNLTVFDKENIHFVPDTYAG-YSTAGADGVIHLVNGRIILKKIQIPDYQRDVTV 82
Bcopc00373     LPAKGNLSLQVFDNANVRFLPNTYPS-FSEADSDGIIHLVNGRIILKKIQIPDYQRDVTV 101
Bfrag0811      LPAKGDLHIPVFENVNVRFSPDTYPDNYNEADGTGVYHLVNGRIILKKITLPEYKRNVSV 85
Bneo101111     LPAKGDQTVRVFEKTNVRFVPGIYPGNYNEADSMGIYHLVNGRIIVKKITLPDYKRNVSV 85
Bzoo00003      HPAMGDLTLRVFDKTPVCFRPDTLKG-YNEPDADGVIRLVNGRIILKKIHLPSYRRNVRV 83
Bhel0045       YPAVGDVNVKVFEKTNVCYRPDRWNG-FNEAGADGVIRLVNGRIILKKIHIPVYKRNVRV 84
EmenII         IWKSRFITIPVFENETINFSDASKSDQ-------VIQRFGNGTIILKKVKIPEIKQGNTI 261
Bhel1482       INQSGVITIPVFDQQPICFNGAKLPAT---LEEGIMYPAGGGTIILKKVKLPEYVKNRSI 264
Bmas03341      INQSGVITIPVFDQQTICFNNAKLPDT---LEDGITYSAGGGTLILKKVKLPESAKNRSI 273
Bdor01199      INQSGVITIPVFDQQSICFNNTKLPEV---LKEGVQYSAGGGTILLKKVKLPDYVKNRTV 271   Group 2
Bvul0565       INQSGVITIPVFDQQSICFNNAKLPEV---LEDGVQYSAGGGTILLKKVKLPDYVKNRTV 271
Bsar02163      INQSGVITIPVFDQQTICFNGAKLPDT---LEEGIQYSAGGGTIILKKIKLPDYVKNRTL 273
Bcopp01102     INQSVVISIPVFSEQRICFNGAKLPEE---VNDQECYSAAGGTIILKNVKLPDYVANRTV 66
Bbar01797      LNQSGVITIPVFNEQTICFNGAKLPDQ---LNDNECYSAAGGTVILKKVKLPEYVANRTV 272
                       :  *.:    :                      .   :*.:          :

EmenI          FIKNECPNKTCDEWDRYANVYVKNK----------------------------------- 70
Blut11613      SAK-VKLTSNGDRWDKSGSCFVIPASSVINMIEVA-AGRAKYPAVDSTKLEHFQGIVPGD 146
Bcopp01256     DIK-VELASNGDRWDKSGSVFVLPKESVINLLNIA-EGKQKFPEVDSTKYENMIGIVPGK 142
Bbar02401      MLH-VQVASNGDRWDKSGSVFVLPKNSPINIMSIA-EGKREFPAIDEARLENMKGIVAGP 142
Bmas01659      KLR-LTLASNGDRWDKSGSCFALPKESFVNLMSIA-QGKAAFPPVDSLKYENMIGIVPGK 147
Bsar01338      TLK-VTVASNGDRWDKSGSCFVLPKESVINLMNIA-EGKKAFPAVDSTKYEKMIGIVPGQ 144
Bvul2763       TLK-VTVASNGDRWDKSGSCFVLPKESVINLMNIA-EGKRAFPAVDSTKYEKMIGIVPGQ 144   Group 1
Bdor04367      TLK-VTVASNGDRWDKSGSCFVLPKESVINLMNIA-EGKRAFPAVDSAKYEKMIGIVPGK 144
Bple01865      SLK-LTVASNGDRWDKSGSCFVLPKNSAVNLLSIA-QGKRKFPEIDSLKLENMIGIVPGK 140
Bcopc00373     SLK-VTVASNGDRWDKSGSCFVLPKNSAVNLLSIA-QGKNKFPEVDSLKLENMVGIIPGK 159
Bfrag0811      SLK-VTLASNGDRWDKSGSCFVLPKSSAINLLTIA-RDGMKFPSVDSLKLEKMVGIVPGK 143
Bneo101111     KLK-LTLASNGDRWDKSGSCFVLPASSAINLLNIA-KGDAKFPKIDSLKLENMNGIIAGK 143
Bzoo00003      AAT-VSVESNGDRWDKTGSCFVLPKESVINMLGVA-RDEQHYPETDTARVELFKGIVSGA 141
Bhel0045       TAT-VTVESNGDRWDKTGSCFVLPRESAVNLLNIA-QGEKRFPAIDSTKLENLKGIIAGD 142
EmenII         FVE-LKQKSNGDAYDRTGDVFIIPQERAISYYTGLTQGVKSLPVYQNGNGKSYQGVALTP 320
Bhel1482       FVE-VAQYSDGDAYDRTGSIFVIPTGKKQSFLDAI-RDLKSVPAFQSK-DMAYPALISTP 321
Bmas03341      FVE-VAQYSDGDAYDRTGSVFVIPTDKKQSFLDAI-RNLKSVPSFQAK-DGNYPALISTD 330
Bdor01199      FAE-VVQYSDGDAYDRTGSVFLIPESKRLSFLDAM-RDLKKVPSFHSE-NMDYHGLISTA 328   Group 2
Bvul0565       FAE-VVQYSDGDAYDRTGSVFLIPEGKQLSFLDAI-RDLKKVPSFRSE-NTDYHGLISTA 328
Bsar02163      FAE-VAQYSDGDAYDRTGSIFMIPTDKAQSFLDAL-RNLKSVPAFQSE-GSDYHGLISTE 330
Bcopp01102     FVE-VAQYSDGDAYDRTGSIFLIPTDKKQSFLDAL-RDLNSVPAFRSG-ETDYHGLVSTD 123
Bbar01797      FVE-VSQYSDGDAYDRTGSVFLIPTDKKQSFLDAL-RNLNSVPSFRSD-STDYHGLVATE 329
                       .  * :*: .. :
```

```
EmenI             TTGEWYEIGRFITPYWVGTEKLPR-------------------GLEIDVTDFKSLLSG-N 110
Blut11613         SFSPNVELMRFMTPFGVGYFSKMDSVTAERRKPVYIDEFAPYAEWEQNITDLYPLLEG-E 205
Bcopp01256        DYLPTVELMRFMTPFGVGHFSAPDDSLSATRRPVYIPHWEKSVTWQQDITQLYPLLEG-E 201
Bbar02401         DYLPTVELMRFMTPFGVGHFSAPDDSLSSKRRPVYIPKWETSVNWVQDITPLYPMLEG-E 201
Bmas01659         DYVPTLELMRFMTPFGVGFYSKKDNEIGAKRKPVYISEWAENVVWEQDITDLYPALEK-E 206
Bsar01338         DYVPTLELMRFMTPFGVGYYSSDNDSLSSKRRPVYIPKWEKSVTWVQDITDLYPALER-E 203
Bvul2763          DYVPTLELMRFMTPFGVGYYSSDNDSLSSKRRPVYIPKWEKSVTWVQDITDLYPALER-E 203
Bdor04367         DYVPTLELMRFMTPFGVGYYSSDNDSLSSKRRPVYIPKWEKSVTWVQDITDLYPALER-E 203
Bple01865         DYLPTIELMRFMTPFGVGHFSEDDDSLSSKRRPVYIPKWEKCVQWEQDITDLYSALKG-E 199
Bcopc00373        NYLPTIELMRFMTPFGVGHYSENDDSLSSKRRPVYIPKWEKCVQWEQDITDLYAALKG-E 218
Bfrag0811         DYLPTVELMRFMTPFGIGHYSNNNDSLSSKRRPVYIPKWESNVTWQQDITDLYPLLEG-E 202
Bneo101111        DYQPTVELMRFMTPFGVGHYSNNEDTLTSKRRPVYIPKWEKQVQWEQDITDLYPLLKG-E 202
Bzoo00003         GYFPAIELMRFMTPFGVGFYSGGDDEKASLRRPVYIDGWAPRAEWTQDVTDRFSSLEG-E 200
Bhel0045          DYLPTLEIMRFMTPFGVGFYSKPDNELSSTRKPVYIDHWEDNVSWTQDVTDRYSALEG-D 201
EmenII            DYLPFIELMRFFTPFGIGHFNEKIQ--------LKGKNWHNNTPYRQDITELRPQLSGKE 372
Bhel1482          YYDTPLELMRFFTAFGVRKFNYN-K--------VKGQDWVDSVLYKSEVTNLAEHLQG-E 371
Bmas03341         DYEAPVELMRFFTGFGVRKFNHN-K--------VKGQHWVDSVIYKSEVTPLASQLQG-E 380
Bdor01199         EYDVPLELMRFFTGFGVRKFNYN-K--------VKGQDWVDSVLYKMEVTPLAEKLEG-E 378
Bvul0565          EYDVPLELMRFFTGFGVRKFNYN-K--------VKGQDWVDSVLYKMEVTPLAEKLEG-E 378
Bsar02163         SYDVPLELMRFFTGFGVRKFNHN-K--------VKGQNWVDSVIYKTEITPLMEKLEG-E 380
Bcopp01102        NYNVPMELMRFFTGFGVRSFNHN-K--------VPGQNWVDSVVYKSEVTPLIERLSG-E 173
Bbar01797         TYDTPIELMRFFTGFGVRSYNHN-K--------VMGQDWVDSVLYKSEVTPLVEHLQG-E 379
                        *: **:* : :   .                          ::*     *.  :
```

Group 1

Group 2

```
EmenI             TELKIYTETWLAKGREYSVDFDIVYGTPDYK---YSAVVPV---VQYNKSSIDGVPYGK- 163
Blut11613         VYVGAFIDTWTKEGYKLSLELDFKESALKCDKLPKRKVVPLVNTVYYY-----GQSIPDL 260
Bcopp01256        AYVGVFIDTWTPEGYVVSMELDIKESKLANDVMPKRHITPLMNTVYYI-----GQTYPDI 256
Bbar02401         AYVGVFIDTWTKEGYLADVRIEVKETEVPCEALPKRQVTPLMNTVYYI-----GQTYPDI 256
Bmas01659         AYIGIFIDTWTAEGYVVGLDIEVKESKISCDALPKRHVQPLINTVYYI-----GQTYPDI 261
Bsar01338         AYVGIYIDTWTAEGYVASMELDVKESKITCDVMPERCVRPLMNTVYYI-----GQTYPDI 258
Bvul2763          AYVGIYIDTWTAEGYVASMELDVKESKITCDVMPERRVKPLMNTVYYI-----GQTYPDI 258
Bdor04367         AYVGIYIDTWTAEGYVASMELDVKESKITCDVMPERRVKPLMNTVYYI-----GQTYPDI 258
Bple01865         VYVGIFIDTWTKEGYVASMELKIKETPVTCEKLVRRHVEPLMNTVYYI-----GQSYPDI 254
Bcopc00373        VYVGIFIDTWTAEGYIASMELNIKETPIACEKLIRRHVEPLMNTVYYI-----GQSYPDI 273
Bfrag0811         AYVGIYIDTWTSEGYLVNADIDVKESRLACDVLPKRHVEPLMNTVYYM-----GQSYPDI 257
Bneo101111        AYVGVFIDTWTPEGYVVSMELDVKESKITCNPLPKRHVEPLMNTVYYI-----GQSYPDI 257
Bzoo00003         AYVGIFIDTWTAEGYVASLTLEVKESAIPEDALLRTRVLPLINTVPYV-----GQSLPDL 255
Bhel0045          VYVGVYIDSWTAEGYLVSLELKVKESDIPEDKLRQTHVLPLVNTVPYQ-----GQNIPDI 256
EmenII            ILIGAFIGNYDKGGHQISLELSIHPD--QQKIVNNNFVLPVFNTTNVMEM--AGQDYPTM 428
Bhel1482          AWIGAYIGNWDAKGHRLSLNLKYYPD--DEHRILK--TIPLFNTVNYLEQ--AGQAYPTF 425
Bmas03341         VWIGAYIGNWDAKGHRLSLKLKYYPD--DERRVNK--AMPLFNTVNYLEQ--AGQAYPVF 434
Bdor01199         AWIGAYIGNWDAKGHRLSLKLKYYPD--EEHRVYN--TLPLFNTVNYLEQ--AGQPYPIF 432
Bvul0565          AWIGAYIGNWDAKGHRLSLKLKYYPD--EEHRVYN--TLPLFNTVNYLEQ--AGQPYPIF 432
Bsar02163         AWIGAYIGNWDAKGHRLSLKLKYYPD--EEHRVYN--AIPLFNTVNYLEQ--AGQPYPIF 434
Bcopp01102        AWIGAYIGNWDAKGHRLSLKLKYYPD--EEHRVYK--SIPLFNTVNYMEQ--AGQPYPIF 227
Bbar01797         AWIGAYIGNWDAKGHRLSLKLKYYPD--DEHRIYK--SIPLFNTVNYMEQ--AGQPYPIF 433
                    :  :  .:   *   .  :.                 *:   .        *
```

Group 1

Group 2

```
EmenI             -AHTLALKKNIQLPTN---TEKAYLRTTISGWGHAKPYDAGSRGCAEWCFRTHTIAINNS 219
Blut11613         FAR-KSLSVPFTLPKNAKNVRLNYITTGHGGHDGGDEF----------VKKENIVSIDGH 309
Bcopp01256        FAR-RAVTTDFTLPRDARNVELKYIVTGHGGHSGGDEF----------VQKQNIVSVDGK 305
Bbar02401         FAR-KPVETTFTLPREAKNVQLKYIVTGHGGHSGGDEF----------VERQNIVSVDGK 305
Bmas01659         FAR-KDVAMDFELPRHAKNVRLKYIVTGHGGHSGGDEF----------VKKRNIVSVDGE 310
Bsar01338         FSR-KDVVMDFDMPEAARNVRLKYIVTGHGGHSGGDEF----------VEKRNIVSVDGK 307
Bvul2763          FSR-KDVVMDFDMPKAAKNVRLKYIVTGHGGHSGGDEF----------VEKRNIVSVDGK 307
Bdor04367         FSR-KDVVMDFDMSKAAKNVRLKYIVTGHGGHSGGDEF----------VEKRNIVSVDGK 307
Bple01865         FAR-KSVSTDFLLPKNAKNVRLKYIVTGHGGHSGGDEF----------VQKRNILSVDGK 303
Bcopc00373        FAR-KSVSADFVLPKNAKNVRLKYIVTGHGGHSGGDEF----------VQKKNILSVDGK 322
Bfrag0811         FAR-RDVSTDFTVPKGAKNIRLKYIVTGHGGHSGGDEF----------VQKRNIISVDGK 306
Bneo101111        FSR-KDVSTEFTVPKGAKNVRLKYVVTGHGGHSGGDEF----------VEKRNILSIDGR 306
Bzoo00003         FAR-RSVTVDAEVPAKAKNLRLQYIATGHGGHSGGDEF----------TQQLNIVRVDGD 304
Bhel0045          FAR-KAVEVPFHLPASARNVRLKYITTGHGGHSGGDEF----------TQQRNLVKVDGE 305
EmenII            FNSDKGVEVEFILTKDLKNAQLRYITTGHGGWGAGDEF----------VPKENSIYLDGK 478
Bhel1482          LGN-DTLRVKFTLNEPVTNARLFYLTTGHGGWGGGDEF----------NQKPNTILLDGQ 474
Bmas03341         FLN-DSLRVRFTLKEPAKNARLFYLTTGHGGWGNGDEF----------NQKPNTVYLDGK 483
Bdor01199         MRQ-DSLIVKFTLKEPAKNARLYYLTTGHGGWGGGDEF----------NQKPNTLYLDGE 481
Bvul0565          MRQ-DSLTVKFTLKEPAKNARLYYLTTGHGGWGGGDEF----------NQKPNTLYLDGE 481
Bsar02163         MRN-DSLTVRFTLKEPVKNARLYYLTTGHGGWGGGDEF----------NQKTNTIYLDGQ 483
Bcopp01102        MLN-DSLKATFTLKEPVKNAKLYYVTTGHGGWGNGDEF----------NQKPNTIYLDGE 276
Bbar01797         MLK-DSLKATFTLKEPVKDAKLYYFTTGHGGWGNGDEF----------NQKPNTIYLDGQ 482
                        :     :       .  *. *  .* . .. :            : : : ::.
```

Group 1

Group 2

Extra twist region

```
EmenI          NTFQHQLGALGCSANPINNQSPGNW---------------------------TPDRAGWC 252
Blut11613      EVIRFIPWRNDCASFRRFNPSTGVWLQKRTASYISEEGKRAEKEIEEPIASSDLSRSNWC 369
Bcopp01256     EALNFIPWRDDCASFRRFNPGTGVWLVKRLASYIDGEGY-AMKEVEEPLGSSDLSRSNWC 364
Bbar02401      TVLDFIPWRDDCASFRRFNPATGVWLIKREAAYIGENGY-EVKEVEEPLASSDLSRSNWC 364
Bmas01659      NVLDFIPWRDDCASFRRFNPGTGVWLIKRLSSYIGKNGY-EEKEIEEPLGSSDLSRSNWC 369
Bsar01338      EVLNFIPWRDDCASFRRFNPATGVWLIPRVAAYIGDKGY-TTKEIEEPLASSDLSRSNWC 366
Bvul2763       EVLNFIPWRDDCASFRRFNPATGVWLIPRVAAYIGDKGY-TTKEIEEPLASSDLSRSNWC 366
Bdor04367      EVLNFIPWRDDCASFRRFNPATGVWLIPRVAAYIGDKGY-TTKEIEEPLASSDLSRSNWC 366  Group 1
Bple01865      EVYSFVPWRDDCASFRRFNPATGVWLIKRLAAYISEDGY-KTKEVEEPLASSDLSRSNWC 362
Bcopc00373     EVYSFIPWRDDCASFRRFNPATGVWLIERLAAYISEDGY-KTKMVEEPLASSDLSRSNWC 381
Bfrag0811      EVLNFIPWRDDCASFRRFNPATGVWLIKRLASYIGEKGY-TEKEVEEPLASSDLSRSNWC 365
Bneo101111     EVMNFIPWRDDCASFRRFNPATGVWLIKRKASYIGEKGY-AEKEVEEPLASSDLSRSNWC 365
Bzoo00003      TVIHFVPWRTDCASFRRFNPTSGVWLVKRKAAYIGEKGY-ETKEIEEALASSDLSRSNWC 363
Bhel0045       TVLDFIPWRTDCASFRRFNPGSGVWLEKRMASFIGKEGY-EMKEIEEPVASSDFSRSNWC 364
EmenII         LAHAFTPWRTDCGSYRLFNPASGNF--------------------EDGLSSSDLSRSNWC 518
Bhel1482       KIITFIPWRDDCGTYRNLNPCSGNF--------------------SNGLSSSDLSRSNWC 514
Bmas03341      KVISFIPWRDDCGTYRNSNPCSGNF--------------------SNGLSSSDLSRSNWC 523
Bdor01199      KVISFVPWRDDCGTYRNWNPCSGNF--------------------SNGLSSSDLSRSNWC 521  Group 2
Bvul0565       KVISFVPWRDDCGTYRNWNPCSGNF--------------------SNGLSSSDLSRSNWC 521
Bsar02163      KIISFIPWRDDCGTYRNWNPCSGNF--------------------SNGLSSSDLSRSNWC 523
Bcopp01102     KVITFIPWRDDCGTYRNWNPCSGSF--------------------SNGLSSSDLSRSNWC 316
Bbar01797      KVITFIPWRDDCGTYRNWNPCSGNF--------------------SNGLSSSDLSRSNWC 522
                   .     .*.:    *   * :                             .*:.**

EmenI          PGMAVPTRIDVLNNSLIGSTFSYEYKFQNWTNNGTNGDAFYAISSFVIAKSNTPISAPVV 312
Blut11613      PGSVVAPYSVLLENLSPGD-HSLNIAIPKAQASEGDKMNHWLVSAYIVWDE--------- 419
Bcopp01256     PGSDVVPETAALGDLKAGT-HTFTVSIPEAQPVKGNELNHWLVSAYLVWEE--------- 414
Bbar02401      PGSDVLPETVELGTLSAGE-HTFKVDIPEAEPVDGDKLNHWLVSAYLVWE---------- 413
Bmas01659      PGSDVVPEEVMLGDLKAGK-HTFKISIPEAQEVDGDKLNHWLISAYLVWDE--------- 419
Bsar01338      PGSDVMPEEAVIGDLAAGK-HSFKVSIPEAQQVMAIN----------------------- 402
Bvul2763       PGSDVMPEEAVIGDLSAGK-HSFKVSIPEAQQVDGNKLNHWLVSAYLVWEE--------- 416
Bdor04367      PGSDVMPEEAVIGDLPAGK-HSFKVSIPEAQQVDGNKLNHWLVSAYLVWEE--------- 416  Group 1
Bple01865      PGSDVLPEVIELPGLQAGK-HTFTVSIPEAQPVNKDELNHWLVSAYLVWEE--------- 412
Bcopc00373     PGSDVVPEEIPLKELQAGK-HTFTVSIPDAQPGNGEELNHWLVSAYLVWDE--------- 431
Bfrag0811      PGSDVVPEEAVIGTLAPGK-HTFTVSIPEAQAVDGNKLNHWLVSAYLVWEE--------- 415
Bneo101111     PGSDVWPEEVALGTLSPGK-HTFSVSIPEAQKIKGNELNHWLVSAYLVWEE--------- 415
Bzoo00003      PGSDVWPVAACLKHLKPGS-HIFAFSIPNAQPAKGEELNHWLISAYLVWEE--------- 413
Bhel0045       PGTDVWPEEVNLSNLVSGS-HILSISVPDAQPADGDKMNHWLISAYLVWDD--------- 414
EmenII         PGTITNPVYINLGNLNAGK-HTIQVKIPQGAPE-GSSQSFWNVSGVLLGQE--------- 567
Bhel1482       PGTVTNPEYICLGNLEAGE-HTLSVQIPQGAPE-GNSNSYWCISGTLIY----------- 561
Bmas03341      PGTVTTPEYIYLGDLEAGE-HTLSVRIPQGAPE-GGSNSYWCISGTLLY----------- 570
Bdor01199      PGTVTNPEYIYLGDLEAGE-HSITVKIPQGAPE-GESNSYWCISGTLIY----------- 568  Group 2
Bvul0565       PGTVTNPEYIYLGDLEAGE-HSITVKIPQGAPE-GGSNSYWCISGTLIY----------- 568
Bsar02163      PGTVTNPEYIYLGDMEAGE-HILSVQIPQGAPE-GGSNSYWCISGTLIF----------- 570
Bcopp01102     PGTVTNPEYIYLGDLDAGT-HTITVKIPQGAPE-GGSNSYWCISGTLIY----------- 363
Bbar01797      PGTVTNPEYIYLGDLEAGT-HTITVQIPQGAPE-GGSNSYWCISGTLIY----------- 569
               **  .      :     *  .     . .

EmenI          TN    314
Blut11613      --    419
Bcopp01256     --    414
Bbar02401      --    413
Bmas01659      --    419
Bsar01338      --    402
Bvul2763       --    416
Bdor04367      --    416                                                      Group 1
Bple01865      --    412
Bcopc00373     --    431
Bfrag0811      --    415
Bneo101111     --    415
Bzoo00003      --    413
Bhel0045       --    414
EmenII         --    567
Bhel1482       --    561
Bmas03341      --    570
Bdor01199      --    568                                                      Group 2
Bvul0565       --    568
Bsar02163      --    570
Bcopp01102     --    363
Bbar01797      --    569
```

**Supplemental Figure S2. Sequence alignment of two PNGases from *Elizabethkingia meningoseptica* and twenty PNGases from thirteen *Bacteroides* species.** A sequence alignment to show key differences between the different PNGases. The N-terminal bowl-like domain found in Group 2 PNGases is highlighted in purple and the extra twist region present in the Group 1 PNGases from *Bacteroides* species. The residues that are key to the specificity of accommodating the α1,3-fucose typical of plant N-glycans are highlighted in the same way as in Fig. 1. The residue blocking the α1,3-fucose in the Group I PNGases is highlighted in blue (E118 in PNGaseF/EMTypeI), the glycine replacing this residue in the Group 2 PNGases is highlighted in pink (G380 in EMTypeII), and the glutamic acid replacing the function of E118 is highlighted in green(E419 in EMTypeII). Alignments were carried out using Clustal Omega (see Methods).

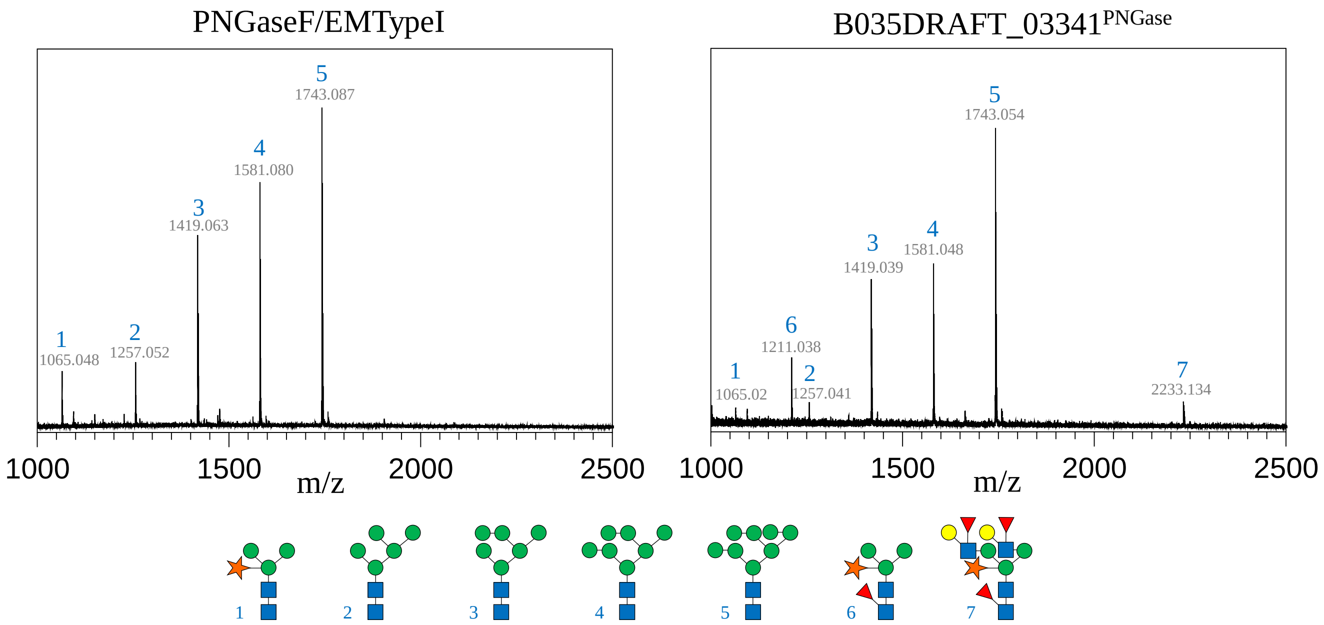


**Supplemental Figure S3. MALDI mass spectra of PNGase activity against soya protein extract.** These data correspond to the samples from Fig. 2E. Prior to labelling, the finished assay was spotted on to a ground steel target on top of Super-DHB matrix. Data was collected using a Bruker Auto-flex Speed in positive ion mode, range 900-3500 m/z at a 50 % laser intensity. Data was processed using Flex analysis 3.5. The data show the presence of plant-type N-glycans 6 and 7 for B035DRAFT_03341$^{PNGase}$, but not with PNGaseF/EMTypeII.

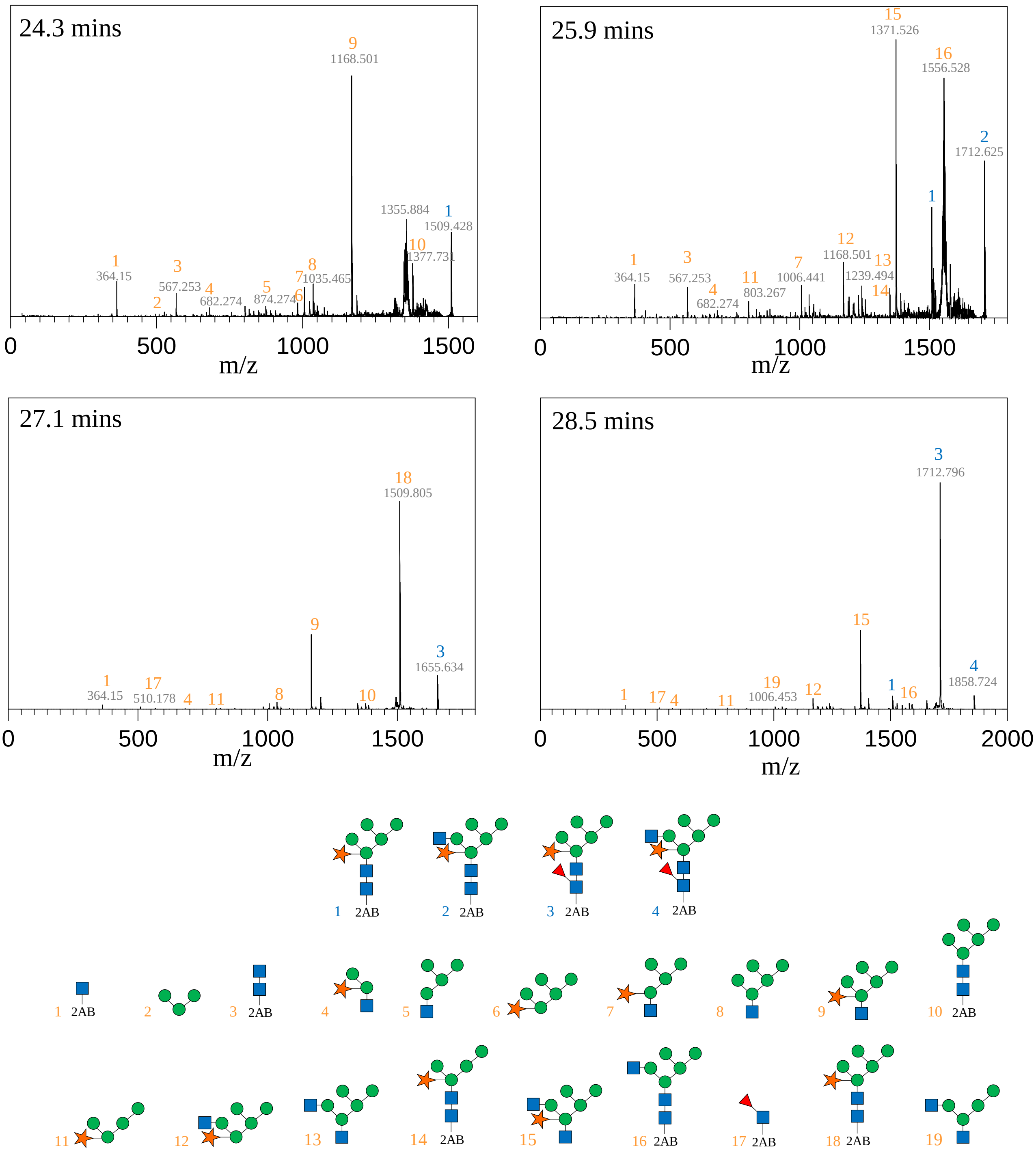


**Supplemental Figure S4. MALDI mass spectra of PNGase activity against papaya protein extract.** These data correspond to the samples from Fig. 2F. The glycans released by B035DRAFT_03341PNGase and labelled with 2AB as described in Materials and Methods. The numbers shown in blue and orange correspond to the full glycan structures and the fragments, respectively.

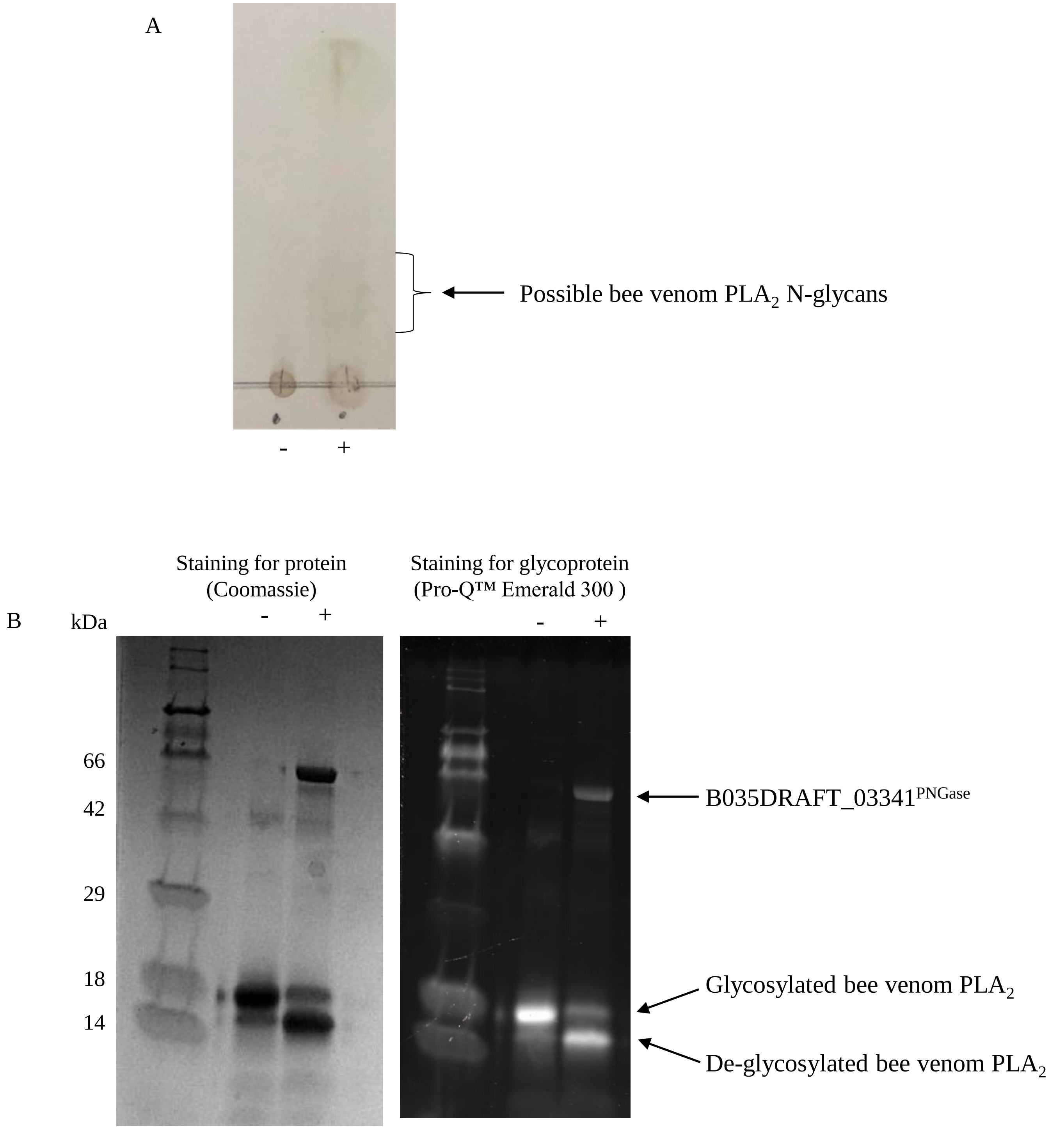


**Supplemental Figure S5. Activity of B035DRAFT_03341$^{PNGase}$ against an insect-derived substrate.** (A) B035DRAFT_03341$^{PNGase}$ was incubated with phospholipase $A_2$ from bee venom to determine if insect N-glycans are also a substrate for this enzyme. Assays contained 1 µM enzyme, 0.5 mg/ml substrate, and 20 mM MOPS pH 7 and were carried out overnight at 37 °C. 12 µl of the assay was spotted onto the TLC plate. (B) The same assay was assessed using SDS-PAGE gels either stained for protein or glycoprotein, where 4 µg of $PLA_2$ was loaded in each lane. A decrease in molecular weight can be seen from the $PLA_2$ in the presence of B035DRAFT_03341$^{PNGase}$ and there is also reduction in glycoprotein staining for this band compared to the control.

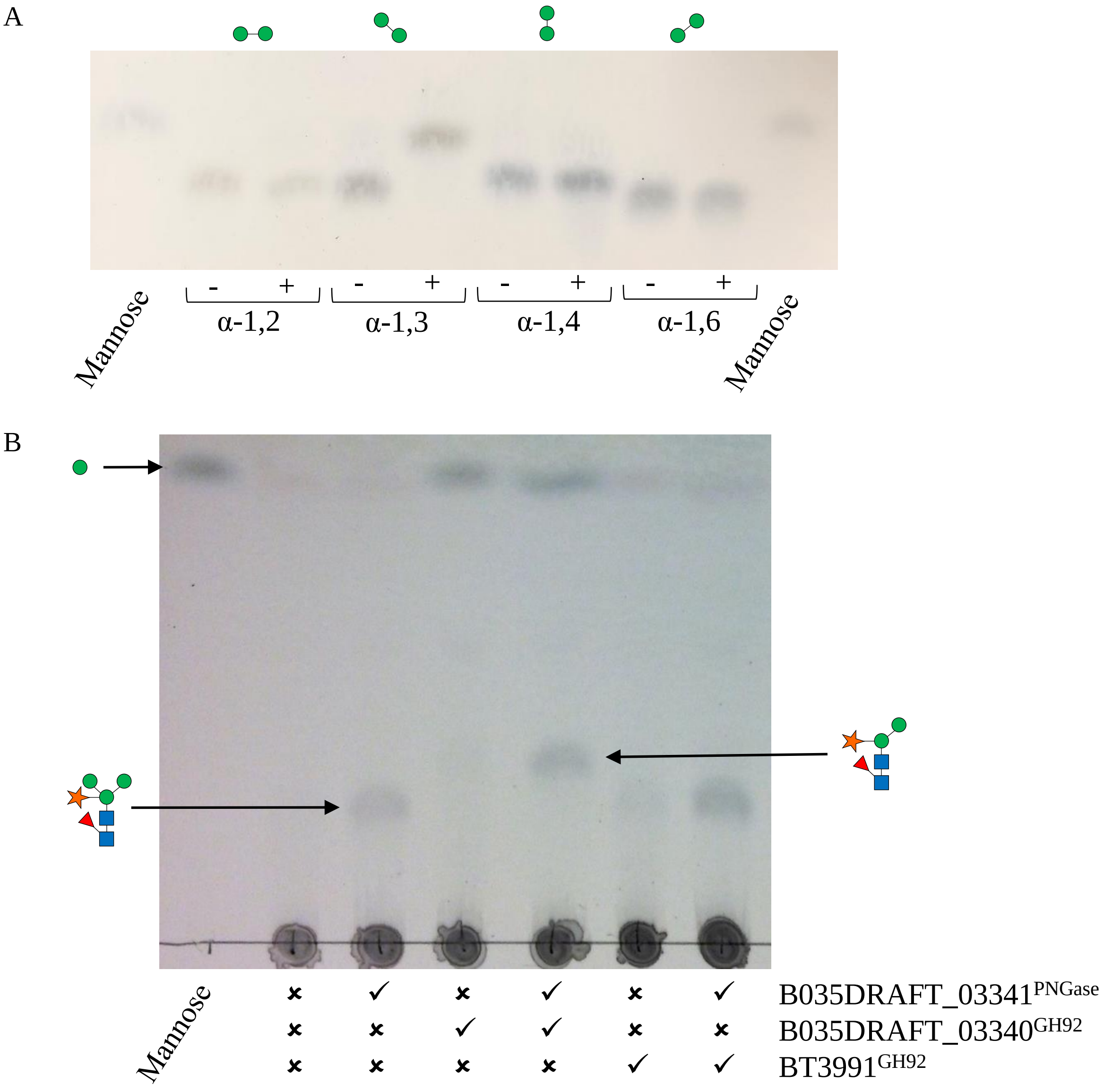


**Supplemental Figure S6. Activity of B035DRAFT_03340$^{GH92}$.** (A) B035DRAFT_03340$^{GH92}$ was incubated with mannobiose with different linkages to determine specificity. Assays contained 1 µM enzyme, 1 mM substrate, and 20 mM MOPS pH 7 and were carried out overnight at 37 °C. 3 µl of the assay was spotted onto the TLC plate. (B) B035DRAFT_03340$^{GH92}$ was incubated against horseradish peroxidase in different combinations with B035DRAFT_03341$^{PNGase}$ and BT3991$^{GH92}$. Assays contains 10 mg/ml of substrate 1 mM substrate, and 20 mM MOPS pH 7 and were carried out overnight at 37 °C. 9 µl of the assay in total (3x 3 µl) was spotted onto the TLC plate.

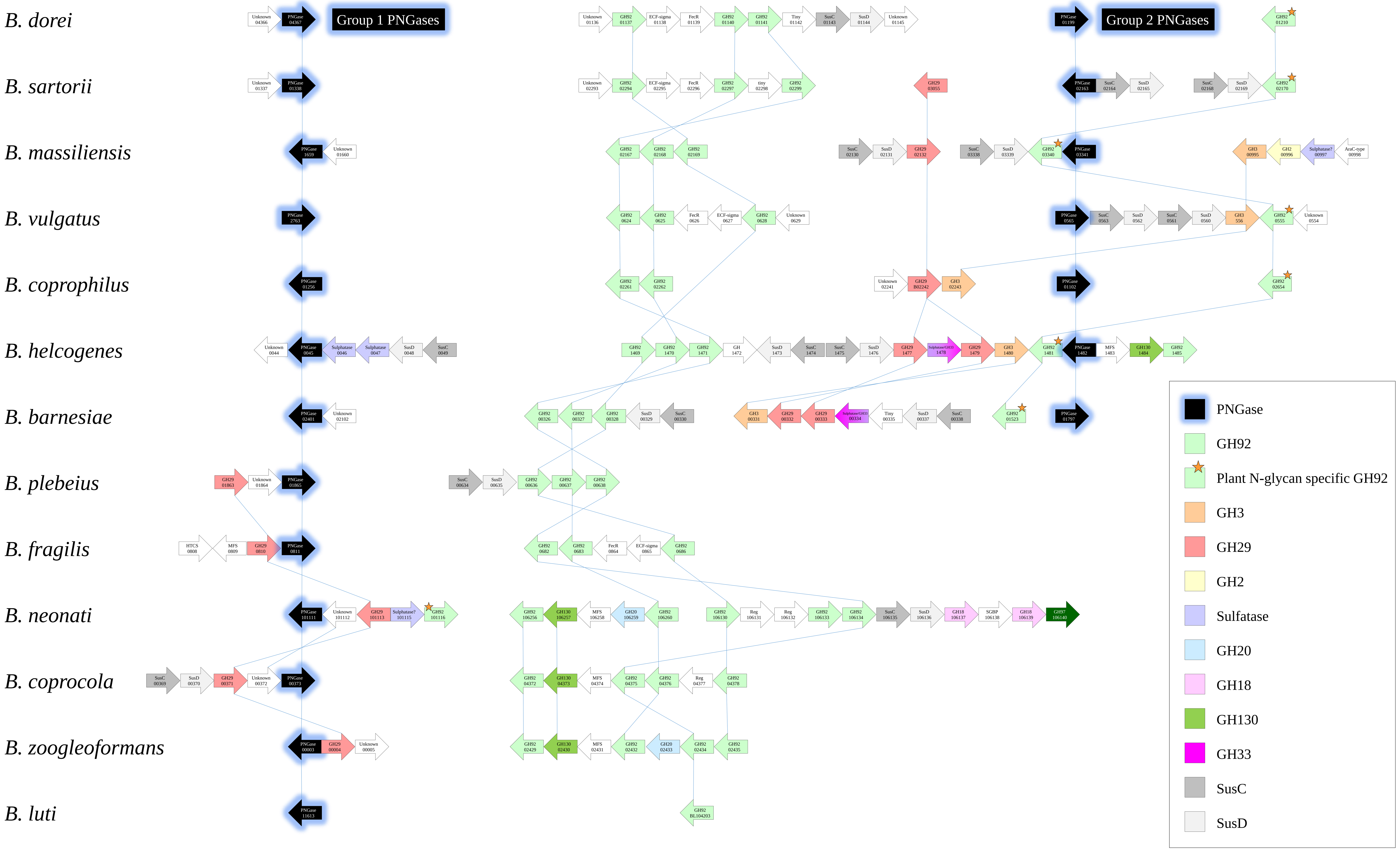


**Supplementary Fig. S7. Functional association analysis of genes associated with PNGase genes in *Bacteroides* species.** The genes in the same loci as the PNGase genes were explored and homologues were identified in the thirteen *Bacteroides* species. The prefix for the locus tags have been left out for brevity, but are as follows: *B. dorei*, *B. sartorii*, *B. massiliensis*, *B. vulgatus*, *B. coprophilus*, *B. helcogenes*, *B. barnesiae*, *B. plebius*, *B. fragilis*, *B. neonati*, *B. coprocola*, *B. zoogleoformans*, and *B. luti* is BACDOR_, JCM17136DRAFT_, B035DRAFT_, BVU_, BACCOPRO_, Bache_, C510DRAFT_, BACPLE_, BF, Ga0057464_, BACCOP_, Ga0052865_, and Ga0131163_, respectively. The blue lines connect the homologues. Abbreviations include: MFS – major facilitator superfamily, Reg - regulation.

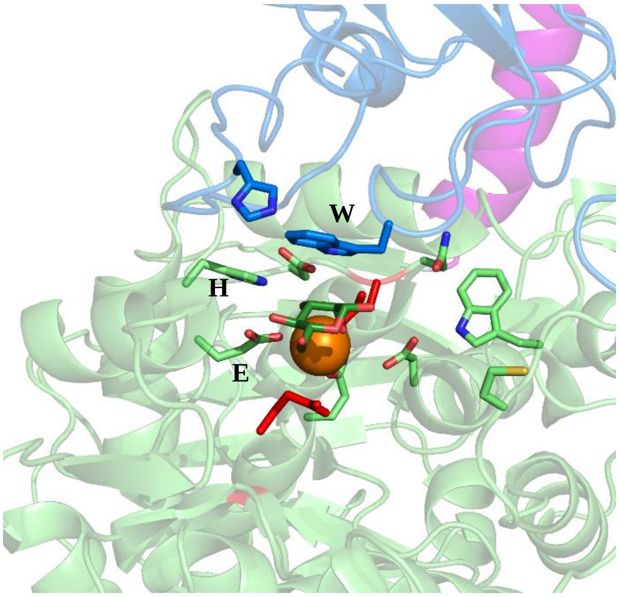


*Streptococcus pneumoniae*
SP2145 5SW1

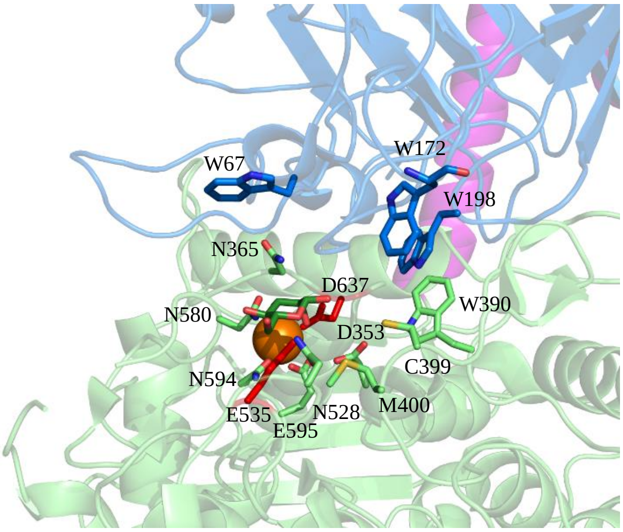


*Bacteroides thetaiotaomicron*
BT3130 6F8Z

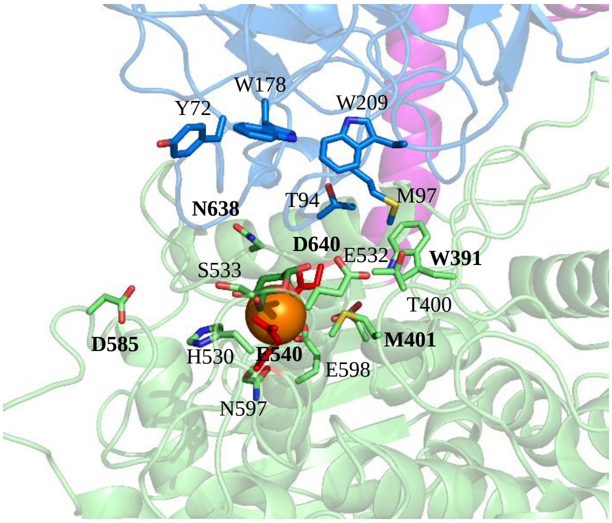


*Bacteroides massiliensis*
B035DRAFT_03340 XXXX

**Supplemental Figure S8. Comparison of the active sites of an α1,2-mannosidase with the two α1,3-mannosidases.** The structure of the α1,2-mannosidase from *Streptococcus pneumoniae* was crystallised with mannose in the +1 subsite (forest green). The three amino acids which drive the specificity for the α1,2-linkage are highlighted. The two available crystal structures for α1,3-mannosidases are also shown with the +1 mannose overlaid into the active site.

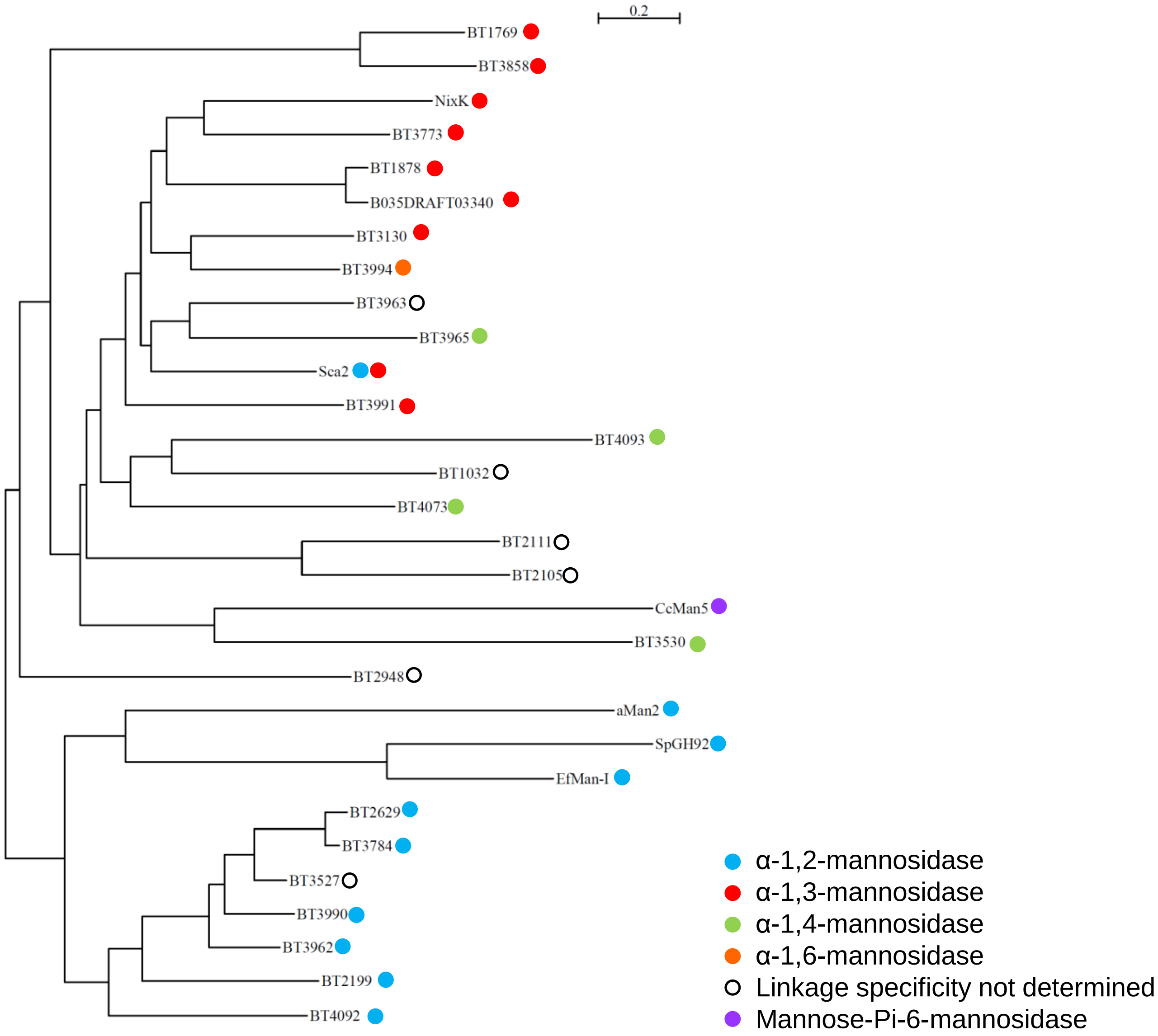


**Supplemental Figure S9. Phylogenetic tree of the characterised enzymes from the GH92 family.** The sequences of GH92 enzymes that have been characterised in the CAZy database were compared as described in Materials and Methods. Specificities can be seen to be grouping although most of the sequences do derive from *B. thetaiotaomicron*.

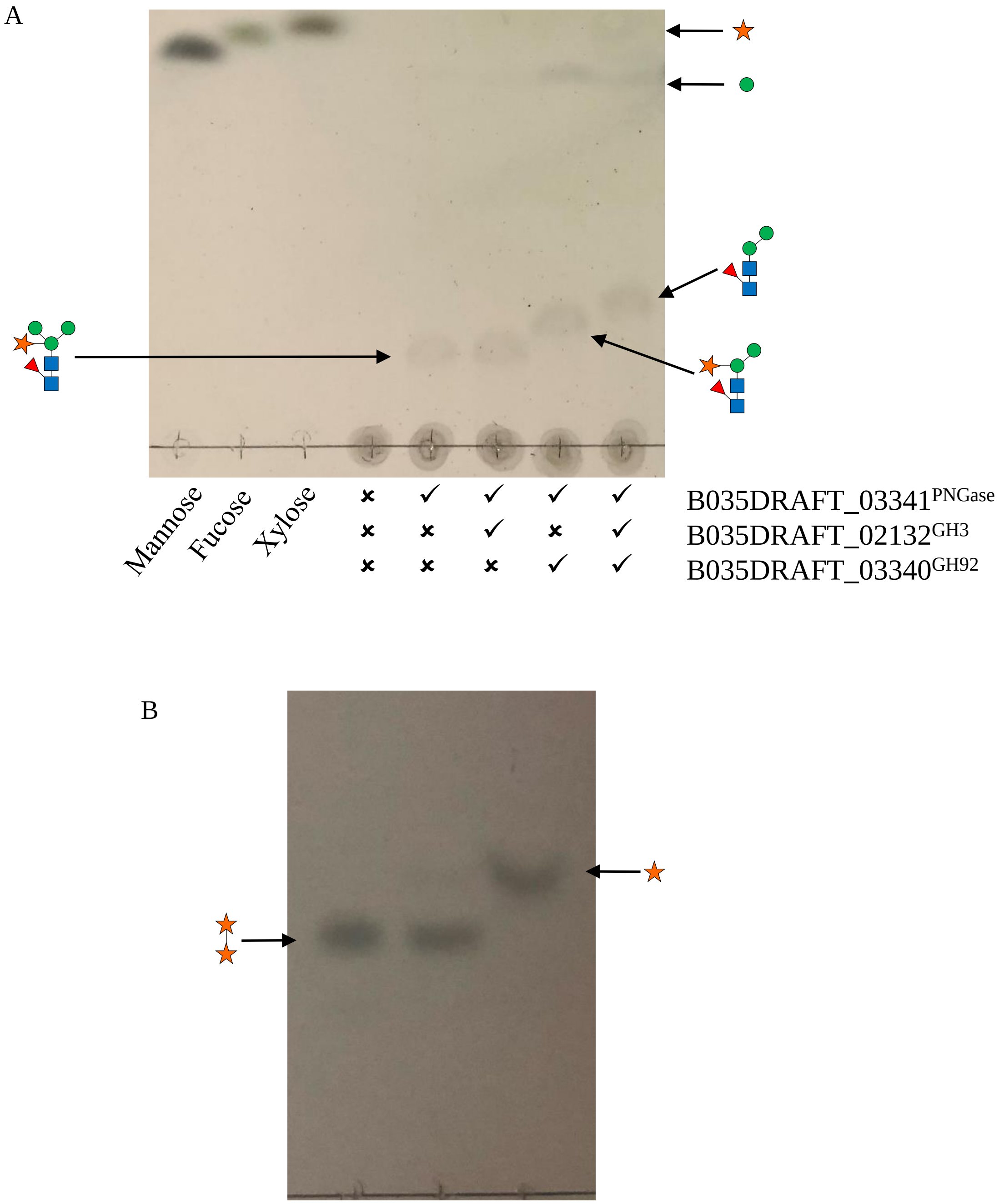


**Supplementary Fig. S10. Activity of B035DRAFT_00995$^{GH3}$.** (A) B035DRAFT_00995$^{GH3}$ was incubated against horseradish peroxidase in different combinations with B035DRAFT_03341$^{PNGase}$ and B035DRAFT_03340$^{GH92}$. Assays contains 10 mg/ml of substrate 1 mM substrate, and 20 mM MOPS pH 7 and were carried out overnight at 37 °C. 9 µl of the assay in total (3x 3 µl) was spotted onto the TLC plate. The results show that B035DRAFT_00995$^{GH3}$ cannot remove the xylose when the α-1,3-mannose is present, but once this mannose is hydrolysed by B035DRAFT_03340$^{GH92}$ then the xylose can be removed also. (B) B035DRAFT_00995$^{GH3}$ was incubated with xylobiose and xylotriose with β-1,4-linkages in comparison to BACOVA_XXX. Assays contained 1 µM enzyme, 1 mM substrate, and 20 mM MOPS pH 7 and were carried out overnight at 37 °C. 3 µl of the assay was spotted onto the TLC plate. (C) BACOVA vs HRP.

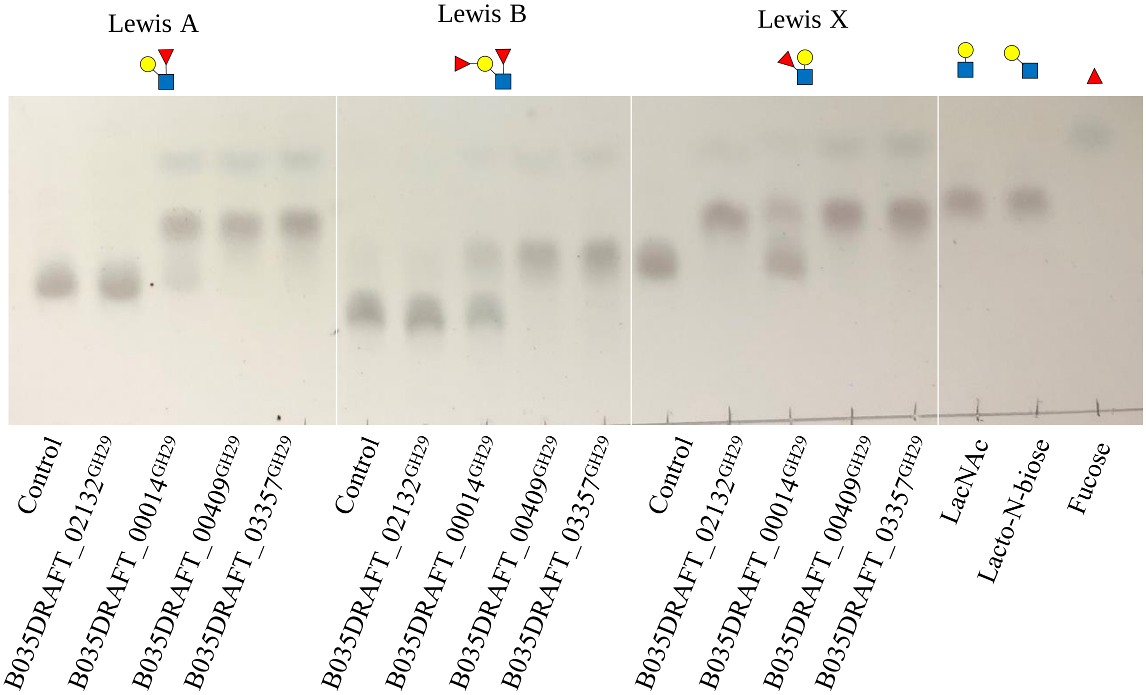


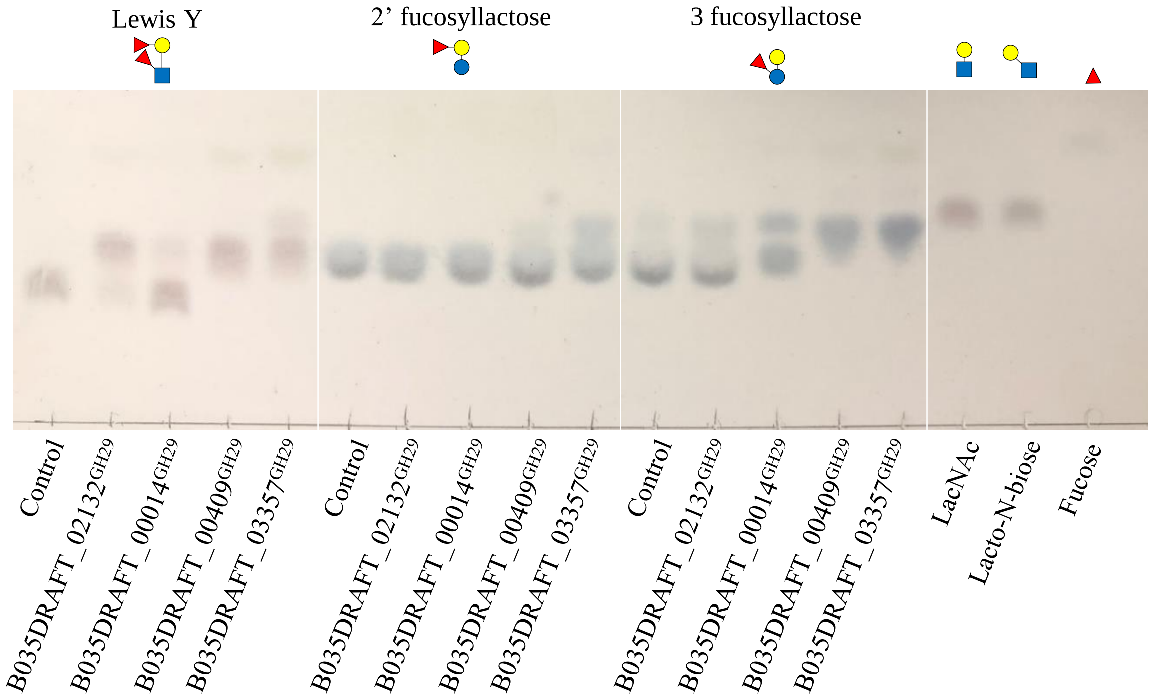


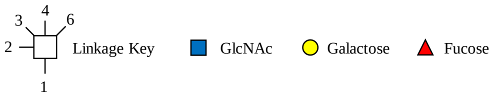


**Supplementary Fig. S11. Activity of four GH29 family members from B. massiliensis.** GH29 enzymes were assayed against different substrates with fucose decorating the non-reducing ends. Lewis A and B is the epitopes for plant and mammalian complex N-glycans, respectively. Assays contained 1 µM enzyme, 1 mM substrate, and 20 mM MOPS pH 7 and were carried out overnight at 37 °C. 3 µl of the assay was spotted onto the TLC plate.

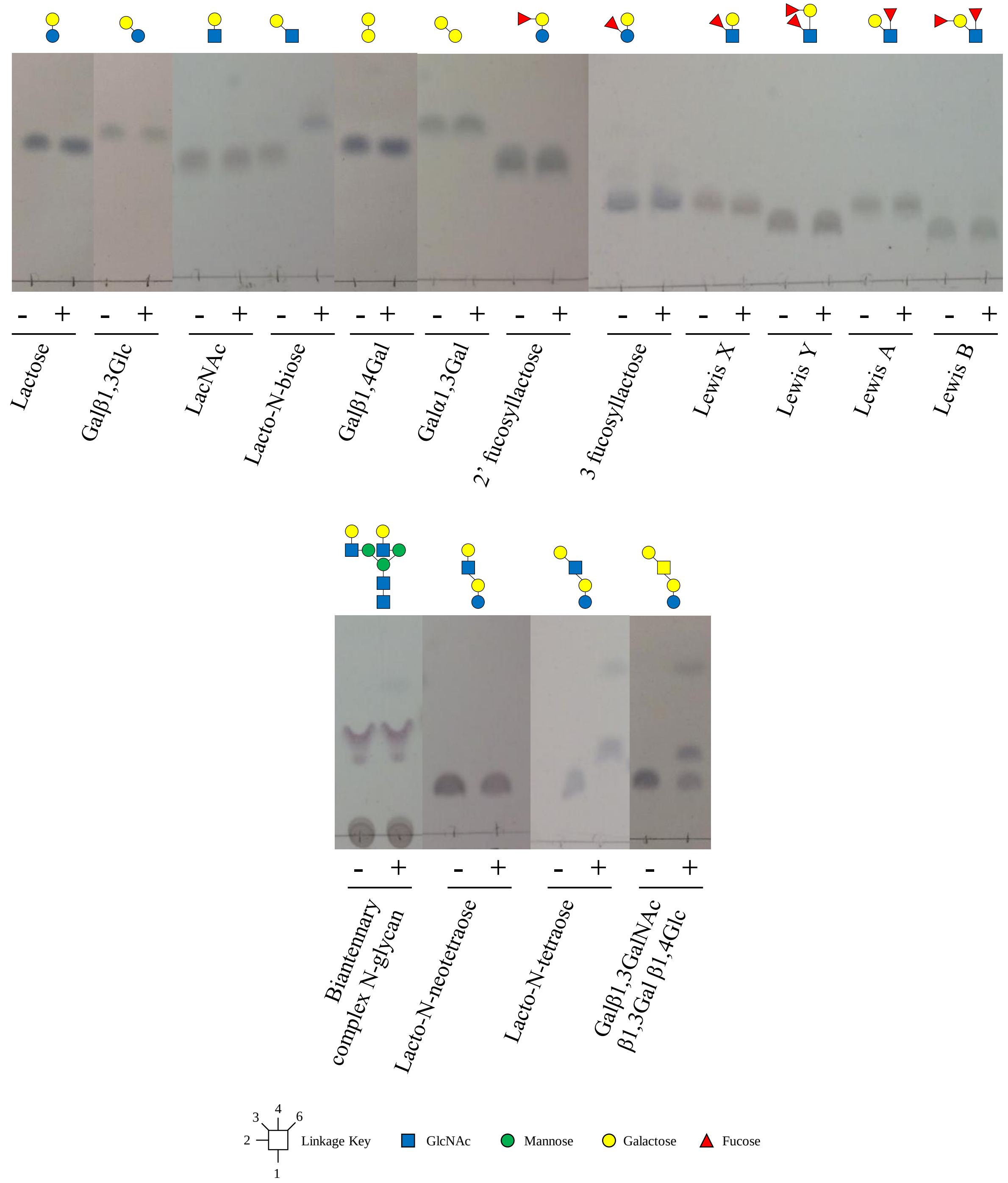


**Supplementary Fig. S12. Activity of B035DRAFT_00996$^{GH2}$ β-1,3-galactosidase.** B035DRAFT_00996$^{GH2}$ was assayed against different substrates with galactose decorating the non-reducing ends. Assays contained 1 μM enzyme, 1 mM substrate, and 20 mM MOPS pH 7 and were carried out overnight at 37 °C. 3 μl of the assay was spotted onto the TLC plate. Only the biantennary complex N-glycan was different with 10 mg/ml substrate and 9 μl of the assay was spotted onto the TLC plate.

A

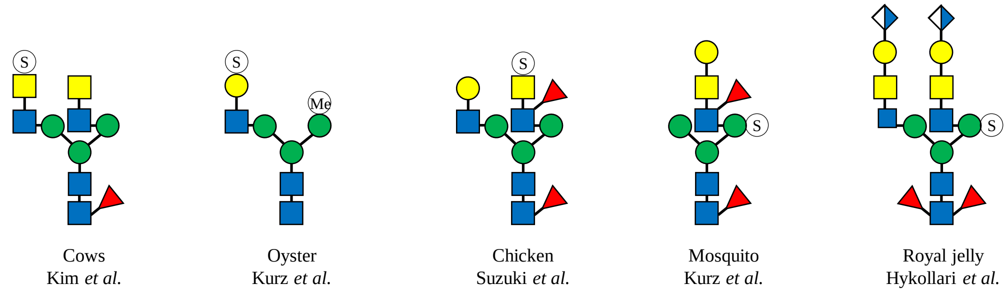


B

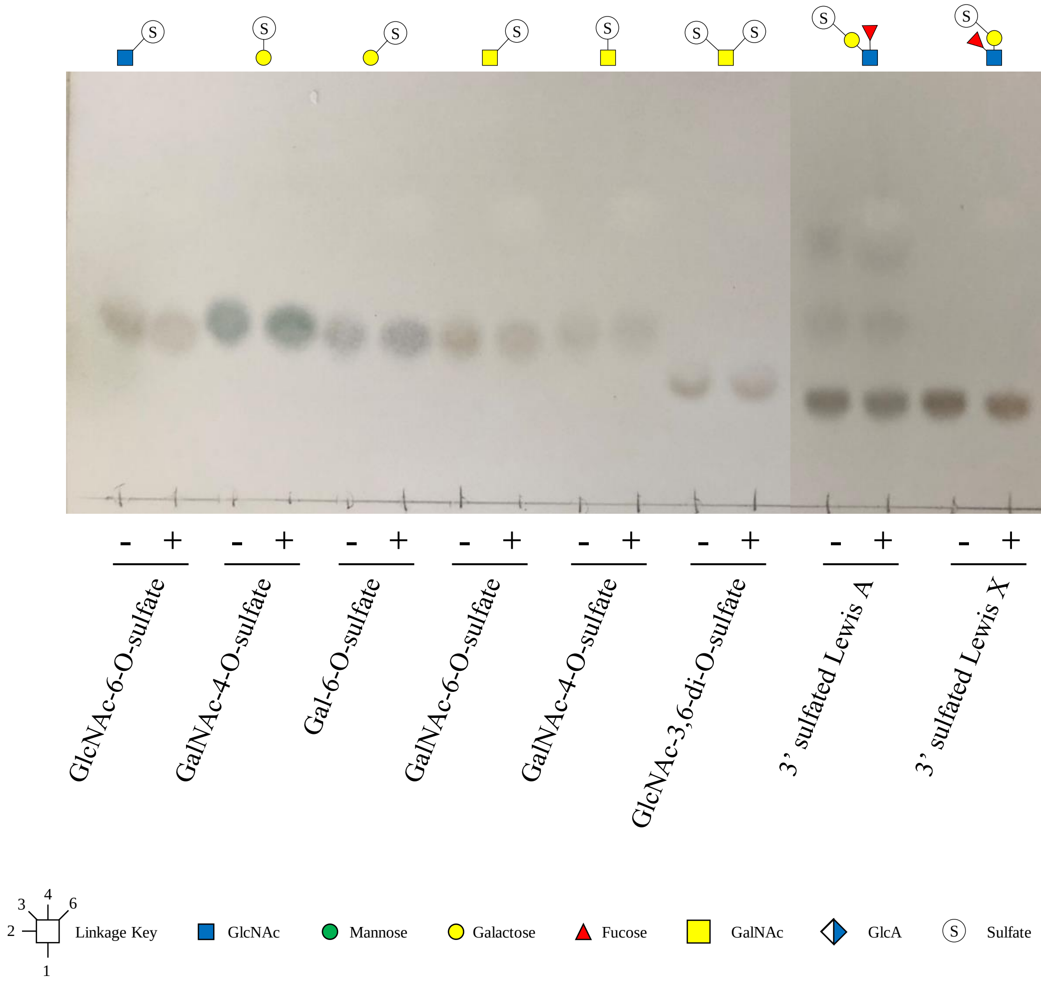


**Supplementary Fig. S13. Sulfated N-glycans and testing the activity of B035DRAFT_00997$^{sulfatase}$** (A) Structures of sulfated N-glycans that have been reported in the literature previously. (B) B035DRAFT_00997$^{sulfatase}$ was assayed against different sulfated substrates. Assays contained 1 µM enzyme, 2 mM substrate, and 20 mM MOPS pH 7 and were carried out overnight at 37 °C. 3 µl of the assay was spotted onto the TLC plate.

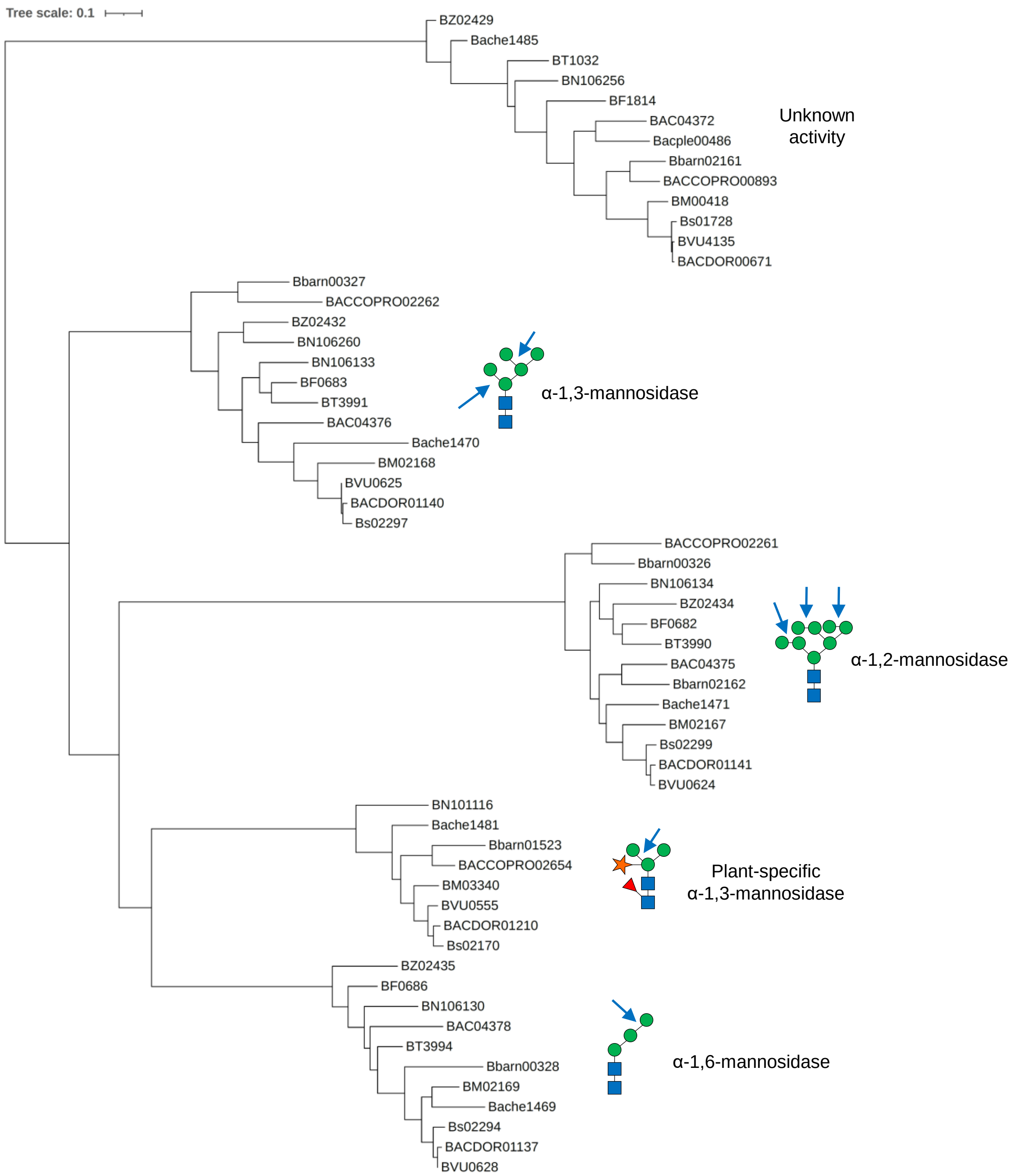


**Supplementary Fig. S14. Phylogenetic tree of GH92 enzymes.** Within the different groups, the GH92 enzymes with unknown function, the α-1,3-specific mannosidases, the α-1,2-specific mannosidases, the plant N-glycan α-1,3-specific mannosidases, and the α-1,6-specific mannosidases have 68-99, 61-99, 70-98, 70-96, and 69-99 % identity between members of that particular branch.
